# DNA polymerase actively and sequentially displaces single-stranded DNA-binding proteins

**DOI:** 10.1101/2025.03.07.642097

**Authors:** Longfu Xu, Shikai Jin, Mia Urem, Seung-Joo Lee, Meindert H. Lamers, Xun Chen, Peter Wolynes, Gijs J. L. Wuite

## Abstract

Single-stranded DNA-binding proteins (SSBs) play a crucial role in stabilizing and protecting transiently exposed single-stranded DNA (ssDNA), yet the mechanisms governing their displacement by DNA polymerase (DNAp) during replication remain largely unexplored. Using bacteriophage T7 DNAp and its SSB, T7 gp2.5, we investigated the molecular mechanisms and visualized the dynamic process underlying SSB displacement. Our single-molecule force spectroscopy demonstrates that T7 SSB modulates DNA replication in an ssDNA conformation-dependent manner, regulated by tension applied to the DNA template. By integrating dual-color single-molecule imaging, we observe that T7 SSB remains stationary as DNAp approaches, indicating that SSB molecules are sequentially displaced rather than pushed forwards. Molecular dynamics (MD) simulations revealed reduced energy barriers for SSB dissociation in the presence of DNAp. This finding, combined with the detected FRET signals when DNAp approaches an SSB-bound ssDNA region and observations of faster replication rates compared to relative slow intrinsic SSB dissociation, collectively support an active displacement mechanism. Using both ensemble and single-molecule analyses, we demonstrated that SSB saturation of ssDNA is critical for optimal replication efficiency, with each SSB molecule contributing positively to the process. Taken together, the uncovered spatial-temporal coordination between SSB and DNAp is necessary for resolving molecular collisions during DNA replication, and may represent a universal strategy employed by other DNA translocating motors to ensure genomic integrity.

## Introduction

Single-stranded DNA (ssDNA) intermediates, arising during biological processes such as DNA replication, recombination, and repair, are susceptible to enzymatic hydrolysis and secondary structure formation(*1*, *2*). These factors could impede DNA polymerase activity and disrupt DNA synthesis(*3*, *4*). In living cells, SSB protein swiftly binds to these susceptible ssDNA intermediates. By potentially stabilizing the ssDNA lagging strand, suppressing DNA hairpin formation(*5*, *6*), and recruiting specific proteins like DNA polymerase (DNAp)(*3*) and helicase(*7*, *8*) to target DNA sites, SSBs are believed to enhance DNA polymerization’s efficiency and processivity during lagging strand synthesis(*3*, *5*, *6*, *9*, *10*). Considering the high affinity of SSBs for ssDNA (micromolar to nanomolar range)(*11–13*), however, the potential for SSB forming molecular roadblocks arises. This, in turn, raises questions about the mechanism through which SSB-DNA complexes are either displaced or advanced along the ssDNA during replication without hindering the replicative polymerase progress, and how resolving these molecular roadblocks carries physiological function.

Single-molecule techniques are powerful tools for studying dynamic biomolecular processes(*14–16*) and have deepened our understanding of replication mechanisms and their regulation. Among the two major types of single-molecule approaches, force-based manipulation studies have revealed that mitochondrial SSBs enhance DNA polymerase γ activity when functional interactions between polymerases and SSBs are firmly established, leading to optimal replication rates(*4*). Concurrently, fluorescence-based visualizations have demonstrated diverse consequences of SSB proteins bound to ssDNA when encountering translocating motor proteins on the same DNA track. For example, human Replication Protein A (RPA) can be displaced or bypassed by helicases like *Xeroderma Pigmentosum Group D*(*17*); *E. coli* SSB can be pushed by a translocase(*18*); and *Pif1*, a helicase from *Saccharomyces cerevisiae*, can even chemo-mechanically propel a human RPA heterotrimer along ssDNA(*19*). However, the specific dynamics of replicative polymerases encountering stationary SSBs, such as T7 SSB(*20*), on the lagging strand, and how SSB binding and secondary structure formation affect replication, remain poorly understood.

Molecular dynamics (MD) simulations can provide atomic-level insights into transient molecular events, challenging to capture with current experimental approaches. The complexity of simulating the polymerase-ssDNA complex stems from its size and the lack of precise force fields for protein-DNA interactions, making modelling this crucial biochemical process rare(*21*). Therefore, MD simulation applications to study the dissociation of SSB and its interactions with partners like DNA polymerases and helicases are limited to a few sample systems, such as human RPA and *E. coli AlkB*(*22*, *23*). Our recently-developed coarse-grained protein model AWSEM has successfully predicted structures and mechanisms in various complex DNA binding systems(*24–26*). Specifically, the benchmarked ssDNA force field from prior work on the simulation of T7 gp4 helicase moving on a long ssDNA has demonstrated AWSEM’s ability to capture the diverse states encountered during translocation’s large-scale motions(*24*).

In this work, we sought to combine these complementary approaches to visualize how DNAp overcomes SSB and resolves molecular collisions on DNA. We first employed an integrated experimental approach, utilizing both force and fluorescence microscopy, to directly monitor the displacement of stationary T7 SSB by replicative T7 DNA polymerase during lagging strand DNA synthesis. T7 SSB interacting with T7 DNA polymerase was selected as the model for investigating the molecular mechanism of the displacement of SSBs by DNA polymerase, due to their well-characterized properties and similarities to mechanisms found in more complex organisms. Our findings revealed that T7 SSB modulates the DNA replication rate dependent on ssDNA template conformation. At 10pN force, when secondary structures can easily form, DNAp accelerates in the presence of SSB. Contrarily, replication decelerates under similar conditions at 20pN, where minimal hairpin formation is expected. Next, using dual-color imaging, we visualized the dynamic interplay between T7 SSB and T7 DNA polymerase, uncovering that SSBs are displaced sequentially during this process. Furthermore, we examined the spatial proximity between DNA polymerase and SSBs, revealing potential direct interaction between the proteins, as evidenced by FRET signals indicating a “collision” event. The MD simulation revealed that the binding energy of the gp2.5-ssDNA complex is reduced in the presence of DNA polymerase, suggesting an active manner of the displacement. Notably, the electrostatic interaction analysis demonstrated the pivotal role of the gp2.5 C-terminal tail in facilitating SSB dissociation by anchoring to the DNA polymerase’s front basic patch region, thereby enhancing displacement efficiency. Experiments with a C-terminal truncated variant of T7 SSB (mut T7 SSB) further validated the critical role of this interaction for efficient replication. Under various forces, our real-time DNA primer extension assays and single-molecule force spectroscopy demonstrated the necessity of saturated SSB and C-terminal-mediated interactions for optimal replication, since the mutant SSB variant displays a reduced efficiency. Collectively, these findings provide a comprehensive mechanistic understanding of how DNA polymerase interacts with SSBs during lagging strand synthesis.

## Results

### Regulation of DNA Replication by SSB Proteins

The binding of T7 SSB to ssDNA has been reported to be dependent on the ssDNA conformation induced by applied force(*13*), leading to the hypothesis that this interaction could influence DNA replication in a force-dependent manner. To investigate how the interplay between tension-regulated secondary structure formation of ssDNA and SSB binding affects DNA replication, we used single-molecule high-resolution optical tweezers to measure DNA length changes, catalyzed by DNA polymerase in the presence of SSB proteins under various applied tensions. In our experimental setup, we tethered a DNA template between two optically trapped beads, enabling precise measurement of T7 DNA polymerase activity under controlled tension and allowing us to explore how varying force levels influence the interaction between DNA polymerase and SSB proteins (**Methods, Figure 1A and 1B**). At a higher tension of 50 pN, the applied template force leads the DNA polymerase enzyme to primarily act in its exonuclease mode, removing nucleotides from dsDNA to generate ssDNA regions, and thus increasing the end-to-end distance (EED) between the two beads(*27*, *28*). This mechanism is effectively utilized to generate ssDNA templates for investigation. Lowering tension to 10 or 20 pN encourages SSB binding to ssDNA and initiates replication by DNA polymerase, displacing SSBs and shortening the EED (**Figure 1B**). Replication trajectories were recorded with and without SSB under these tensions (**Figure 1C**), providing insights into the dynamic interplay between secondary structure formation, SSB binding, and DNA polymerase replication activities.

**Figure 1.**
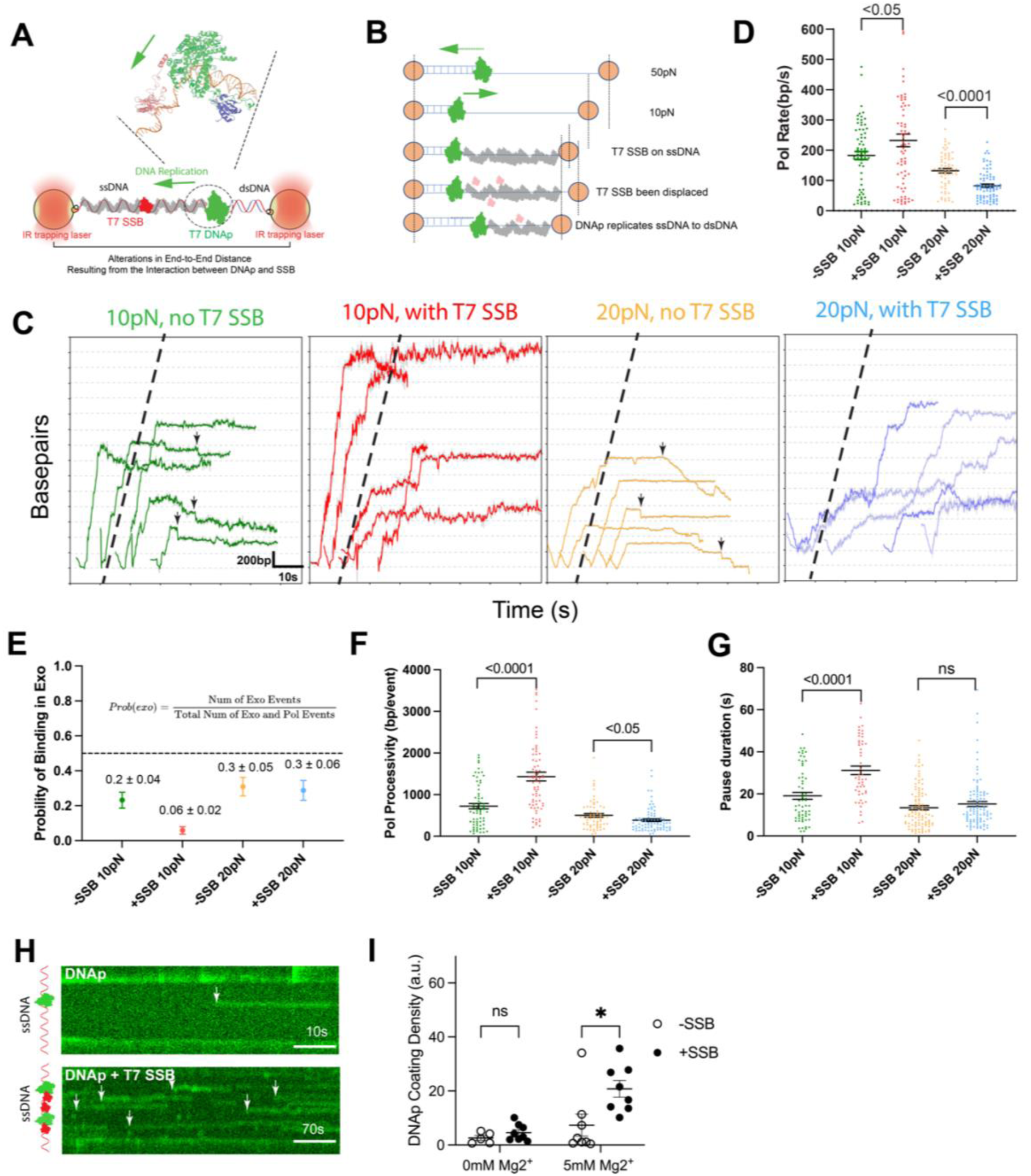
Single-Molecule Analysis of SSB’s Effect on DNA Replication. **(A)** Schematic representation of the experimental setup. The diagram depicts a partially ssDNA and dsDNA template tethered between two optically trapped beads using dual-optical tweezers. The DNA template (∼8kbp) features a 25 nt overhang for T7 DNA polymerase loading. Initially, a high tension of 45pN/50 pN is applied to digest approximately 5 kbp of dsDNA, creating a lengthy ssDNA section as a template. The tension is then reduced to 10-20 pN to analyze polymerization activity in the presence or absence of SSB. The DNA polymerase activity is investigated by observing the alteration in DNA extension caused by the interplay between DNAp and SSB. T7 DNAp (30 nM) and T7 gp2.5 (150 nM) are suspended in a buffer containing 20 mM Tris-HCl (pH 7.5), 100 mM NaCl, 5 mM MgCl_2_, 1 mM DTT, and 0.02% BSA. The magnified inset illustrates the T7 DNA polymerase (PDB ID: 1T7P) and T7 gp2.5 (SSB) (PDB ID: 1JE5) at the ssDNA/dsDNA junction, with distinct colors indicating their identity. **(B)** Diagram illustrating changes in end-to-end distance due to DNAp activity, SSB binding and DNAp and SSB interactions under varying tensions, as measured by high-resolution optical tweezers. At 50 pN, the DNA polymerase exhibits exonuclease activity, removing nucleotides and increasing end-to-end distance. As tension decreases to 10 or 20 pN, the reduced force shortens end-to-end distance, further accentuated by SSB binding to ssDNA. Lowered tension causes DNA polymerase to reverse direction, initiating replication, displacing SSB, and lengthening end-to-end distance. The transition from ssDNA (modelled as FJC^34^) to double-stranded DNA (modelled as WLC^35^) enables activity quantification in the presence of SSB. End-to-end distance traces over time are used to calculate the percentage of ssDNA and changes in base pairs over time. Shifts in polymerase or exonuclease activity are determined using a change-point detection algorithm. **(C)** Representative replication trajectories with and without SSB at 10 pN and 20 pN. Dashed black lines indicate the DNAp replication rate at 10 pN without SSB as a reference. Backtracking dynamics are evident in DNA replication without SSB (indicated by black arrows). Note that base pair-time trajectories include 3-5 seconds prior transition to tension of 10 or 20 pN and are aligned based on the time of tension transition; backtracking dynamics due to high tension at 50 pN are not emphasized. **(D)** Illustrating the effect of SSB on replication efficiency at 10 pN and 20 pN. The changepoint detection approach computes the most likely change points within single-molecule base pair-time traces (**Method**). At 10 pN, SSB considerably enhances replication efficiency from 183 ± 13 bp/s (N=67) to 232 ± 21 bp/s (N=72), with a *p*-value of 0.0467. Conversely, at 20 pN, SSB reduces efficiency from 132 ± 7 bp/s (N=63) to 83 ± 5 bp/s (N=81), with a *p*-value < 0.0001. Data is represented as mean ± standard error of the mean (SEM). **(E)** Displaying the relative probability of DNA polymerase binding to its exonuclease active site with or without SSB at 10 pN and 20 pN. The binding probability in exo was determined by dividing the number of exo events by the total number of exo and polymerase (pol) events for a given DNA molecule. Without SSB, DNAp exhibits a higher probability of exo binding at 20 pN (0.3 ± 0.05, N=30) than at 10 pN (0.2 ± 0.04, N=35), indicating tension-dependent binding to the exo site. With SSB, binding probabilities at the exo site decrease to 0.06 ± 0.02 (N=47) and 0.3 ± 0.06 (N=43) at 10 pN and 20 pN, respectively. All scenarios display a preference for binding at the replication site, with exo binding probabilities less than 50%. Data is represented as mean ± SEM. **(F)** Illustration of SSB’s modulation of replication processivity under varying DNA template tensions. Processivity, measured by the total base pairs synthesized by DNAp per burst event, increases from 722 ± 63 bp/event (N=67) to 1433 ± 104 bp/event (N=72) at 10 pN in the presence of SSB, with a *p*-value < 0.0001, while decreasing from 503 ± 43 bp/event (N=63) to 386 ± 31 bp/event (N=81) at 20 pN, with a *p*-value of 0.0312. Data is represented as mean ± SEM. **(G)** Depicting average pause duration with or without SSB at 10 pN and 20 pN. In the absence of SSB, pause duration decreases from 19 ± 2 s (10pN, N=61) to 13 ± 1 s (20pN, N=101), possibly due to a hairpin structure at 10 pN impeding replication. With SSB, pausing increases to 31 ± 2 s at 10 pN (N=50) but remains relatively constant at 15 ± 1 s at 20 pN (N=99). Data is represented as mean ± SEM; sample size are the same with panel D. **(H)** Representative kymographs showing fluorescently labeled DNAp (30 nM DNAp-Snap-surface 549, ∼60% labeling efficiency) binding to ssDNA substrate in the absence or presence of SSB. Only the green channel is displayed for clarity. White arrows highlight selected DNAp trajectories. **(I)** Analysis of DNAp coating density on ssDNA with or without SSB in buffer containing 0 mM or 5 mM Mg^2+^. Coating density is the normalized difference between the average photon count per pixel in the data window and the background. N = 5-8 DNA molecules. With 5 mM Mg^2+^, the presence of SSB increases DNAp binding to ssDNA (*p*<0.05).

We analyzed the EED changes using a change-point detection algorithm to identify transition points in base-pair time traces, which marks shifts into either polymerase or exonuclease activity under 10pN or 20pN (see **Methods**). Template tensions of 10 pN and 20 pN were chosen as example force thresholds based on their distinct influences on ssDNA conformation and potential effect on SSB-ssDNA interactions. At 10pN, conducive to secondary structure formation, the role of SSB proteins is anticipated to be critical. Our data (**Figure 1D**), derived from analyses of 30-47 different DNA molecules, indicate that at 10 pN, SSB proteins enhance the replication rate from 180 ± 10 bp/s (mean ± SEM, N=67) to 230 ± 20 bp/s (N=72). This increase of approximately 20% is statistically significant with a *p-value* of 0.05, confirming previous biochemistry studies under no template tension which show that SSB proteins bolster DNA replication by precluding the formation of secondary structures(*5*, *6*, *10*). Intriguingly, at an increased tension of 20 pN, surpassing the threshold for secondary structure disruption and reducing the likelihood of their formation, the presence of SSB proteins led to a decrease in replication efficiency from 130 ± 10 bp/s (N=63) to 80 ± 5 bp/s (N=81) (mean ± SEM, *p*-value < 0.0001). This suggests that when there is a diminished necessity for SSB proteins (where secondary structures are less prone to form), their presence might impede replication.

Apart from the SSB’s impact on replication rate, our data also indicate a role of SSB proteins in modulating the binding site preference of DNA polymerases. This preference was quantified by calculating the ratio of exonuclease (exo) events to the sum of exo and polymerase (pol) events for a given replication event (**Figure 1E**). Notably, the absence of SSB proteins led to increased backtracking dynamics in DNA replication (black arrows, **Figure 1C**), with a higher probability of DNA polymerase binding at the exonuclease site, particularly at 10 pN (∼20%) and 20 pN (∼30%). In the presence of SSB proteins, the probability of binding at the exonuclease site was reduced to approximately 10% at 10 pN but was unchanged at 20 pN (∼30%). Across all conditions, a preference for binding at the replication site was observed, with probabilities of binding at the exonuclease site consistently below 50%, supporting the notion that lower tension (<∼35pN) favours replication activity (**Figure 1B**) (*27*, *28*). These findings imply that SSB proteins, by binding to the ssDNA in front of the replication fork, may directly interact with the replicative DNA polymerase, thereby promoting its affinity for the replication site.

Moreover, the binding site preference of DNAp directly influences replication processivity (**Figure 1F**). At a tension of 10 pN, the presence of SSB proteins markedly enhanced processivity, increasing from 720 ± 60 bp/event (N=67) to 1,400 ± 100 bp/event (N=72). This enhancement suggests that SSB proteins contribute to replication processivity by both suppressing secondary structures and promoting binding at the replication site. We note that, the processivity values reported here represent apparent processivity, as rapid exchange of DNAp at the replication fork(*29*) was not considered. Interestingly, when the tension was elevated to 20 pN, the inclusion of SSB proteins led to a decline in replication processivity by around 30%. This can be attributed to the lower replication rate (**Figure 1D**) and relatively high probability of exonuclease binding (∼20%) and high exonuclease rate (**Figure 1E, S1A**) at this tension, which may together interfere with efficient replication.

We further analyzed the average duration of pauses both in the presence and absence of SSB proteins (**Methods, Figure 1G**). Without SSB proteins, we observed a decrease in average pause duration from 19 ± 2 s at 10 pN (N=61) to 13 ± 1 s at 20 pN (N=101). At the lower tension of 10 pN, the replication process is likely more impeded by hairpin structures. Such structures which are more prevalent at 10pN might stall DNAp, leading to longer pauses. Surprisingly, with SSB proteins, pause durations increased to 31 ± 2 s at 10 pN (N=50), while at 20 pN the effect of SSB on pause durations was relatively minor (15 ± 1 s, N=99). The pronounced increase in pause duration at 10 pN suggests that while SSB proteins play a crucial role in facilitating replication by preventing secondary structures, they may also introduce significant roadblocks under certain conditions. DNA polymerase was earlier reported to bind to the ssDNA region(*29*), creating self-imposed roadblocks impeding replication. We employed fluorescently labeled DNA polymerase to monitor its binding dynamics to ssDNA in the presence or absence of SSB proteins (**Figure 1H**). Increased binding events of DNAp to ssDNA was observed in the presence of SSB proteins than that in the absence of SSB (white arrow, **Figure 1H**), especially in the presence of 5 mM magnesium ions (our experimental conditions, **Methods**, **Figure 1H** & **1I,** N=5-8 DNA molecules, *p*<0.05). These observations suggest that SSB proteins on ssDNA may facilitate the recruitment of DNA polymerase to the replication fork, but excessive recruitment could lead to self-obstruction and prolonged pause durations.

### DNA Polymerase Displaces Stationary SSBs in a Sequential Manner

During replication, bound SSBs must be displaced by the replicative DNA polymerase to prepare DNA synthesis (**Figure 2A**). In this study, we propose two distinct hypotheses to explain the displacement process: the sequential “one-by-one” model, where there is a gradual eviction of SSBs from the ssDNA, and the “whole-train” model, where there is a simultaneous pushing multiple SSBs in a row. To evaluate these hypotheses, we employed single-molecule dual-color imaging techniques in combination with optical tweezers, facilitating observation and quantification of the interaction dynamics between T7 DNA polymerase and T7 SSB proteins at a single-molecule resolution. Our experimental design features a fully saturated SSB concentration with a small proportion of labelled SSB (∼4%) to distinguish between the displacement models. The fluorescently labelled SSBs served as markers to determine the relative distance between the replicative DNA polymerase and the SSB proteins. For the “one-by-one” model, we anticipated a progressive decrease in distance between the labelled SSB and DNAp, while for the “whole-train” model, we expected that the distance would remain constant over time (**Figure 2A**).

**Figure 2.**
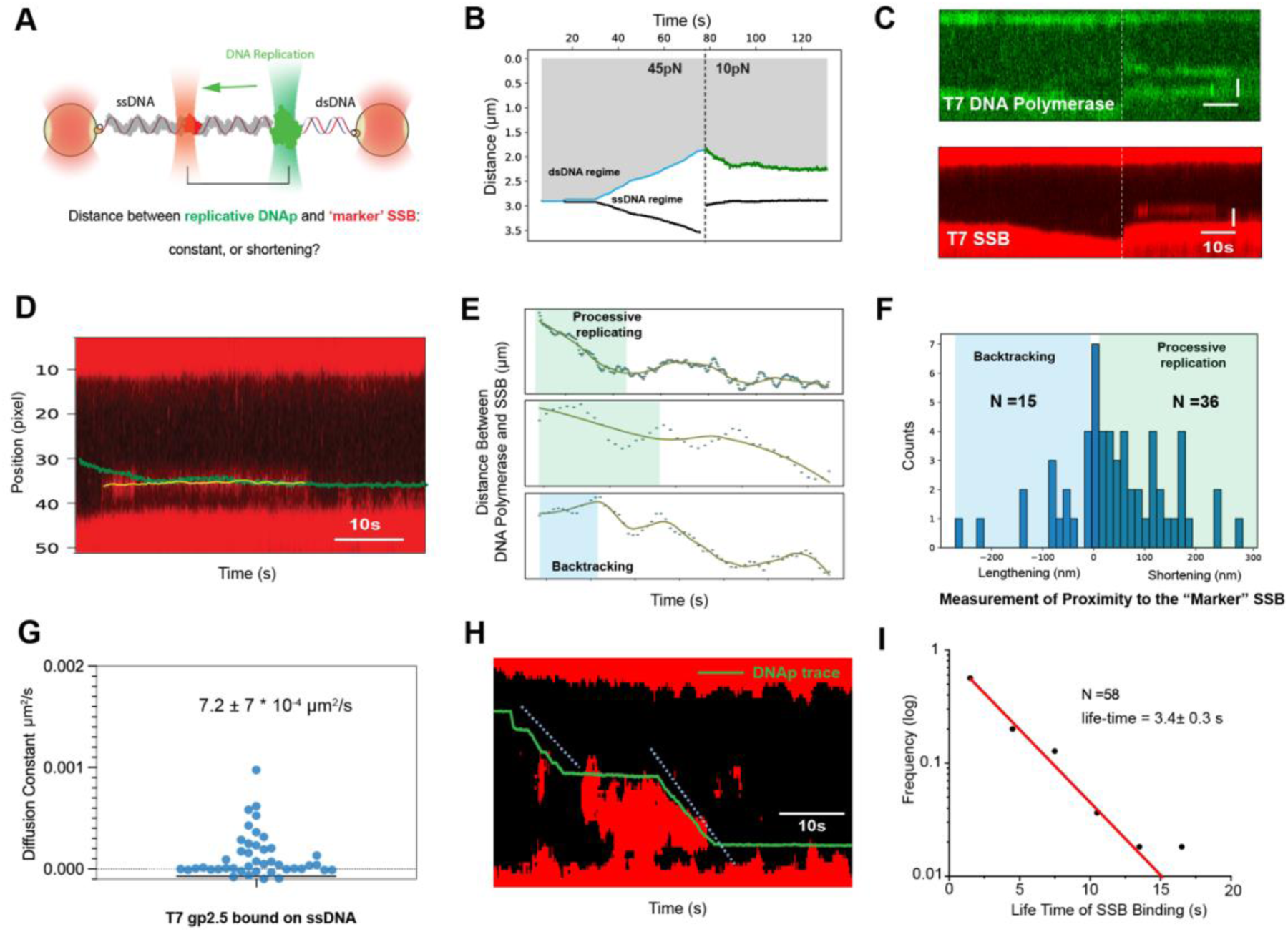
Single-Molecule Filming of T7 DNAP Displaces T7 SSB protein. **(A)** Schematic illustrating the experimental setup to investigate the displacement of SSB by DNA polymerase during replication. A dual-optical trapping system holds a partially ssDNA/dsDNA pkyb1 DNA construct, enabling real-time measurements of end-to-end distance (EED) between two beads. Simultaneously, a dual-color imaging system visualizes fluorescently labeled T7 DNA polymerase (T7 DNAp-snap-surface 549) and T7 SSB (T7 SSB-Atto647N). A strategic concentration of fully saturated SSB with partially labelled SSB (150nM, with an average 4% labelling efficiency), is employed to guarantee persistent interaction with DNAp during DNA synthesis. This setup enables the testing of two models for SSB displacement: the one-by-one-SSB-removal model and the pushing-the-whole-train model. **(B)** Representative mechanical trace illustrating DNAp and SSB interactions; the dashed line denotes the transition from 45 pN to 10 pN force. The calculated ssDNA/dsDNA junction position and EED are plotted against time. The junction position is derived from independent optical tweezer measurements, capitalizing on elasticity differences between ssDNA and dsDNA, to track DNAp in real time. **(C)** Concurrent confocal kymographs for the molecule depicted in **(B)**, capturing dynamic interactions between fluorescently labeled DNAp (top panel) and SSB (bottom panel). The yellow dashed line signifies the force transition. Time scale: 10 s; position scale: 75 nm per pixel. **(D)** Composite image combining data from **(B)** and **(C)** to directly observe DNAp and SSB interactions. Here only the replication process is displayed for clarity (**Figure S4A**). Fluorescently labeled SSB is visualized binding to ssDNA using fluorescence microscopy. SSB trajectories are extracted for further analysis employing a trajectory analysis method (**Methods**). Only SSB trajectories lasting over 3 pixels (approx. 1s) are analyzed. In total, we extracted 58 DNAp-SSB interaction traces from 25 unique DNA molecules (out of 64 DNA molecules) which exhibited clear and consistent fluorescence signals. **(E)** Three presentative distance dynamics between DNA polymerase and SSB during DNA replication using the method **(D)**; the top panel is based on data from panel **(D)**. The distance between DNAp and SSB decreases during processive replication (highlighted in light green), interspersed with DNAp backtracking (highlighted in light blue) and pausing events (highlighted in light gray). Note, the time axis for these three examples is rescaled for presentation purpose. **(F)** Histogram displaying changes in distances between replicative DNAp and SSB during SSB displacement. Only the first burst segment is analyzed to account for potential DNAp forward and backward movements. Negative values indicate backtracking events. Of the 58 events analyzed, 15 exhibit DNAp moving away due to backtracking activity, 36 indicate DNAp moving closer through SSB displacement, and seven events remain stationary, likely attributed to DNAp pausing. These measurements support the one-by-one-SSB-removal for SSB displacement by DNAp. **(G)** Diffusion constant for T7 SSB is computed using the Mean Square Displacement (MSD) method, revealing that SSB predominantly remains stationary on ssDNA. Approximately 10% of the diffusion constants with negative values, possibly obscured by noise, are excluded from the analysis. **(H)** Representative traces showing the removal of multiple closely positioned fluorescent SSB molecules during DNA replication. Be noted that the SSB kymograph was processed with median filtering and thresholding to enhance contrast for better visualization. The DNAp trajectory (green line) is determined using the method explained in panel **(B)**. Time scale: 10 s; position scale: 75 nm per pixel. **(I)** Quantification of average SSB lifetime, determined by fitting a mono-exponential decay to the extracted trajectory durations, yielding an average binding duration of 3.4 ± 0.3 s (N=58).

We executed high-resolution mechanical measurements and aligned those data with fluorescence images of DNA polymerase and SSB, to track the relative movement of DNAp and SSB in real-time (**Figure 2B-2D**). By making use of the differential elasticity between ssDNA and double-stranded DNA (dsDNA), and calibrating the shortening effect of SSB binding to ssDNA (**Methods**, **Figure S2A**)(*13*), we can trace the real-time movement of the ssDNA/dsDNA junction, the site where replicative DNAp binds and catalyses the conversion of ssDNA into dsDNA (**Figure 2B**). The solid blue and green lines in **Figure 2B** denote examples of backtracking (exo) and replicative (pol) activity, respectively. Note that when we change the template tension from 45pN to 10pN, DNAp replicates ssDNA into dsDNA(*27*, *28*). Next, we obtained confocal microscopy images of fluorescent DNAp and SSB simultaneously by separating the channels and rectifying signal crosstalk (**Figure 2C**). The SSB kymograph was investigated with a trajectory analysis method (see **Methods**) for additional quantification (shown in yellow, **Figure 2D**). Subsequently, we superposed the data acquired from DNAp’s position (**Figure 2B**) onto the SSB kymograph image to directly track both the DNAp and SSB positions (**Figure 2D**, **S3**). Using this quantitative approach, we can determine the change in distance between replicative DNAp and SSB (**Figure 2E**). One should note that due to the potential of DNAp moving toward SSB then reversing (as for example the lightblue area in **Figure 2E**), only the first segment with replication activity was quantified. We observed for a majority of events (36 out of 58 traces) that DNAp was moving closer to and displacing SSB (**Figure 2F**). In 7 out of 58 traces, the DNAp appeared to be stationary in the first segment of the trace, potentially due to pausing. Finally, in approximately a quarter of molecules (15 out of 58 events) we observed DNAp moving away from the SSB, attributable to backtracking. Importantly, in none of the traces did we see the SSB move directionally in sync with the DNAp. In fact, the SSB remains stationary on the ssDNA template while the DNAp is moving as can be determined by the very low diffusion constant (7 ± 7 * 10^-4^ μm^2^/s (mean ± SEM; **Figure 2G**) of T7 SSB, calculated using the Mean Square Displacement (MSD) method(*13*). By comparison, the known static protein EcoRV has an apparent diffusion constant of 1*10^-4^ µm^2^/s(*30*). Taking all these observations together, the sequencial one-by-one model seems to explain the data best. And indeed, we can occasionally observe DNAp directly removing indivisual SSBs (**Figure 2H** as examples, **Figure S4**, N= 2 out of 25 DNA molecules).

To determine whether the removal of SSB is a consequence of the active motion of DNAp or the passive dissociation of SSB, we compared the intrinsic dissociation rate of SSB with the DNA replication rate. By fitting a mono-exponential decay to the binding durations observed in the kymographs, we calculated an average SSB lifetime of 3.4 ± 0.3 s after correcting for photobleaching (**Figure 2I**), consistent with prior reports(*13*). This corresponds to an intrinsic off-rate (Koff) of approximately 0.3 s⁻^1^. The measured replication rate is approximately 200 nt/s (**Figure 1D**), allowing DNAp to translocate a distance equivalent to the footprint of SSB (∼10 nt) in 0.05 s. This translates to a potential displacement of ∼20 SSB/s, a rate roughly two orders of magnitude faster than the spontaneous dissociation rate of SSB from ssDNA. This disparity suggests that SSB is likely being actively displaced by the advancing DNAp during replication rather than passively dissociating. If SSBs were to spontaneously dissociate at such rapid rates, it would compromise their protective role, especially during DNAp’s proofreading activities that generate extensive regions of ssDNA. Therefore, our findings support the hypothesis of active displacement of SSB by DNAp, likely involving direct physical interactions between the two proteins.

### Active Displacement of SSB by DNA Polymerase

The high replication rate of DNAp (**Figure 1D**) poses challenges to experimentally capture the structural dynamics of the interaction between DNAp and SSB during replication in real-time. To investigate the structural dynamics in the event of the DNAp-induced SSB displacement from ssDNA, we employed AWSEM MD simulations to explore the intermediate states. T7 SSB has a total of 232 amino acids with a conserved oligosaccharide-oligonucleotide binding fold (OB-fold) that consists of a five-stranded anti-parallel β barrel capped by an α-helix on one end (αA)(*31*). Since the most flexible C-terminal region (residue 206 - 232) and the linker region (residue 79 - 88) are missing in the current available template (PDB ID:1JE5), we modelled the full-length structure using Alphafold2(*32*) (**Figure 3A**). The C-terminal is positioned in close contact with the positive binding groove in gp2.5 in the Alphafold2 model. The ability to predict residue contacts with state-of-the-art accuracy using Alphafold2 provides us with a promising inference of the apo state of gp2.5 without DNA.

**Figure 3.**
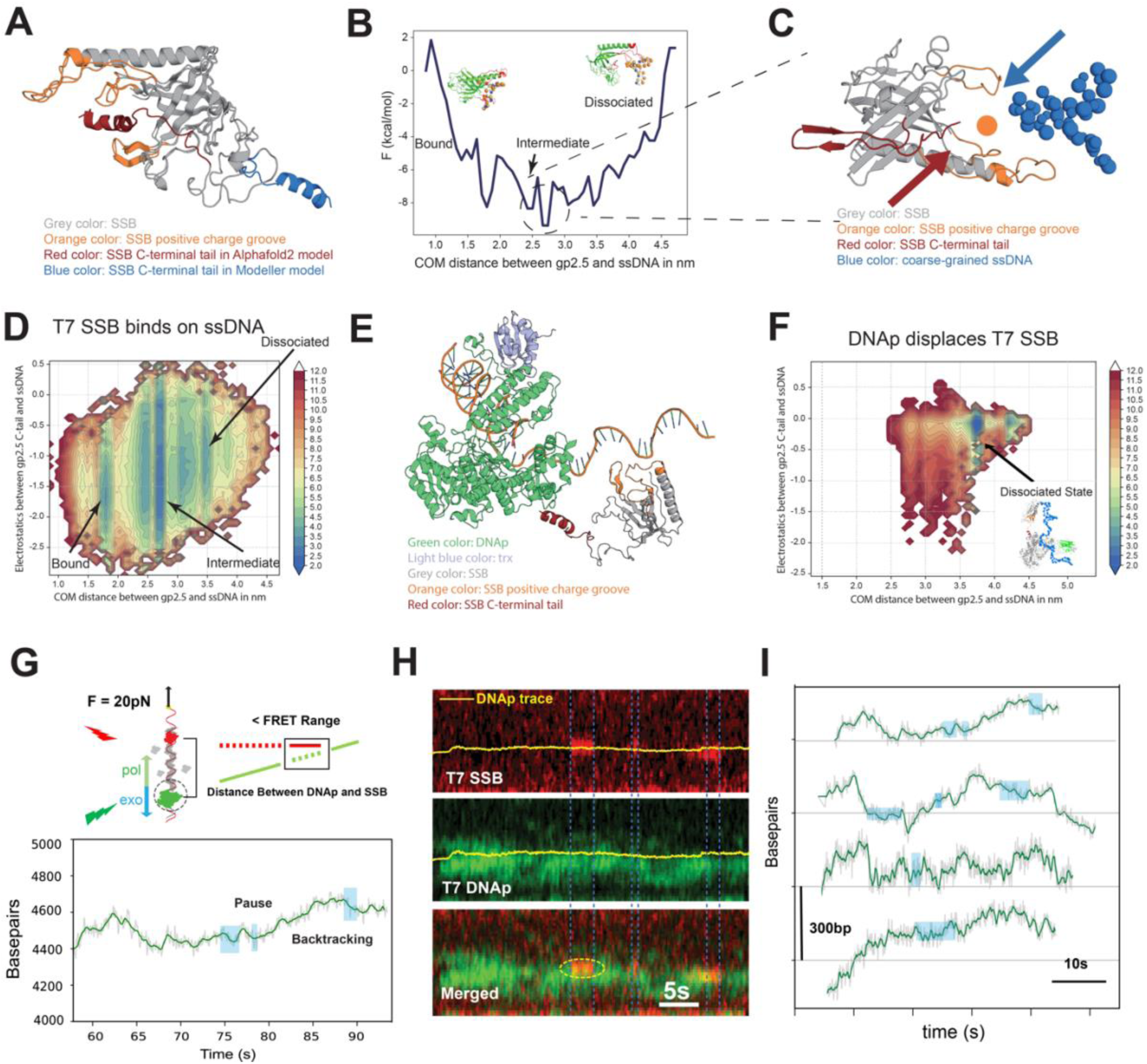
Active displacement of SSB by DNA polymerase possibly by direct physical interaction. **(A)** The structural details of the predicted full-length SSB monomer structures. The majority of the AlphaFold2 predicted structures and Modeller predicted structures colored in grey are the same as the template 1JE5, as expected. Orange color shows the positive binding groove of SSB. Red color shows the C-terminal tail predicted by AlphaFold2, while blue color shows the C-terminal tail predicted by Modeller. **(B)** The free energy profile for the process of SSB binding to the ssDNA. The free energy curve shows three local minima during the binding process. **(C)** The representative structure of the intermediate state in **(C)** is shown. The ssDNA (blue) and C-terminal tail (red) are located in the opposite directions of the positive binding groove, indicating a competitive scenario between them. **(D)** The two-dimensional free energy profile of the SSB binding to the ssDNA. The representative structures of each local basin are also shown. **(E)** The structural details of the DNAp-trx-SSB complex binds to DNA with predicted SSB structures from AlphaFold2. The SSB is positioned based on the template provided in 6P7E. Green color shows the DNAp full-length structure. Blue color shows the thioredoxin (trx) structure. Same as Figure 3A, the grey, orange, and red color show the majority, positive binding groove, and C-tail of SSB. The dsDNA with a 5’ ssDNA overhang is also shown. **(F)** The two-dimensional free energy profile of the SSB binding to the ssDNA with the existence of DNAp and trx. The representative structure of the dissociated state is shown. The binding process is tentative to dissociate with the high electrostatic effect triggered by the C-terminal of SSB. **(G)** Top: Schematics explains the setup for examining the proximity between molecules during SSB displacement by DNAp. Bottom: Representative replication activity data from mechanical measurements, displaying changes in the number of base pairs over time. Notably, regions with relatively constant base pairs, indicating pausing events, and reductions in base pairs, indicating backtracking activity, are observed. These regions correlate with the FRET signal region highlighted in light blue shadow area (from panel **H**). **(H)** Representative kymograph displaying the dynamic interaction between fluorescently labeled DNA polymerase and SSB from the same molecule of panel **(G)**. The upper and lower panels respectively show the fluorescent trajectories of T7 SSB (top panel) and T7 DNAp (middle panel) under confocal microscopy. Only the green laser is used for excitation to visualize the fluorescently labeled DNAp. The kymographs are processed to reduce background noise, correct for signal leakage (crosstalk), and are filtered for clarity. The yellow line represents the real-time position of DNAp calculated from mechanical measurements. Bottom panel shows a merged kymograph from T7 DNAp and T7 SSB. **(I)** We analysed 8 individual FRET events induced (red) SSB signals. These were selected from an initial set of 16 DNA molecules based on signal quality of their fluorescence signals. The corresponding base pair-time traces are extracted, and the region with the FRET signal is highlighted in light blue. Interestingly, in the detected FRET signal regions, pausing events are observed in 6 out of 8 cases and backtracking events in 2 out of 8 cases.

We first explored the SSB binding dynamics on ssDNA. Umbrella sampling was used to systematically sample the conformational space, facilitating the subsequent analysis of the free energy landscape. The one-dimensional (1D) free energy profile revealed three distinct states characterizing the SSB-ssDNA binding process **(Figure 3B)**, leveraging the center of mass (COM) distance between the protein and ssDNA as the primary collective variable. In the initial bound state, this distance is at 1.3-2.0 nm, while the distance in the intermediate state is fluctuated around 2.7 nm, and finally increase to 3.5-4.0 nm in the dissociated state. The hierarchically clustered heatmap and dendrogram indicate two main clusters for the bound state structures (**Figure S5A** and **Figure S5B**). Further examination of these structures unveiled two predominant forms of the C-terminal tail, characterized by disorder and an anti-parallel beta-sheet conformation (**Figure S5C** and **Figure S5D**). Notably, as the COM distance between SSB and ssDNA increased, competition arose between the C-terminal region and the ssDNA for occupancy within the positively charged binding groove, leading to the stabilization of an intermediate state. We extracted the representative structures of the three largest clusters from the cluster map of all structures in the local basin of intermediate state (**Figure S6**). From these structures, we observed the ssDNA was displaced by the C-terminal of the SSB **(Figure 3C)**. The ssDNA shifted from the center of the binding groove toward the one side of the SSB while the C-terminal of SSB attacks from the other side and occupied the binding groove, indicate a transition of the binding process. A morphing movie among those three representative structures suggests a possible trajectory (**Movie S1**). We also extended the 1D profile to a two-dimensional one with an additional axis showing the strength of electrostatics interactions between positive binding groove and ssDNA (**Figure 3D**). Notably, the observed energy change of approximately 5 kcal/mol that the SSB needs to pass to reach the dissociated state from the bound state in the 2D free energy profile aligns with the experimentally derived average binding free energy of -4.7 kcal/mol (**Table S1**). This simulation agrees well with the recent report that SSB remains essentially immobile on ssDNA rather than diffusing with frequent hops on and off the ssDNA(*20*). Next, we computed the 2D free energy profile for T7 SSB dissociation induced by DNAp during replication. The structure of the full length T7 SSB predicted from Alphafold2 was assembled with the DNAp-primer/template DNA complex (PDB ID: 6P7E) with a short peptide that mimics the C-terminal of other components(*33*) (**Figure 3E**). To characterize how the interaction between gp2.5 C-terminus and DNAp facilitates the gp2.5 dissociation from the bound ssDNA, we performed an umbrella sampling simulation to explore the energy difference among local states during the stripping process **(Figure 3F)**. By comparing the 2D free energy profile with that obtained from T7 SSB alone (**Figure 3D**), we observed that the bound state becomes relatively unstable in the presence of DNAp. The free energy profile indicates that the complex quickly becomes trapped in the dissociated state, possibly due to the interaction between DNAp and SSB. From the representative structures of the dissociated state, we observed that the SSB was stripped from ssDNA while maintaining its connection with DNAp through the C-terminal region (**Figure 3F**). This suggests that DNAp likely facilitates SSB removal in an active manner by altering the free energy required for SSB displacement from ssDNA via the C-terminus of SSB.

This active displacement model revealed by MD simulation predicts there is a possible direct physical contact between DNAp and SSB. Indeed, direct contacts between the SSB C-terminal and DNAp, as well as the SSB positive binding groove and ssDNA are observed in some of the representative structures from the movie snapshots (**Figure S7**). Considering the FRET detection range (< ∼10 nm) (*34*, *35*) and the size of T7 SSB and T7 DNAp, estimated to be around 4 nm(*31*) and 7 nm(*36*), respectively, this physical contact could result in a possible FRET signal. Therefore, we aimed to experimentally search for the FRET signals using single-molecule FRET and precise force measurements. If the active displacement hypothesis holds, we would anticipate detecting FRET signals indicative of direct interaction as DNAp displaces SSB during replication. The rapid replication rate that should result in up to 20 SSBs being displaced per second coupled with the relatively low temporal resolution of confocal scanning (approximately 0.7s per scan along the DNA template), however, posed challenges in precisely discerning FRET signals during the course of processive replication. Recognizing these constraints, we exploited the strengths of this approach by measuring the spatial proximity of these proteins during DNA replication under a tension of 20 pN. This force was chosen because DNA polymerase is more likely to backtrack under such conditions (**Figure 1E**), yet T7 SSB still effectively binds to ssDNA(*20*). Higher tension results in reduced T7 SSB binding, while lower tension decreases backtracking likelihood. With our chosen force, the fluorescently labeled DNAp can synthesize dsDNA and occasionally backtrack (**Figure 1C** and **1E**), creating free ssDNA on the DNA template for SSB to rebind from solution. These backtracking events with subsequent SSB rebinding and potential for pausing thus provide valuable conditions to have both physical proteins very close on the DNA, allowing us to measure the physical distance between DNAp and SSB.

Our approach used the fluorescently labelled DNAp-SNAP-Surface® 549 (green) approaching the fluorescently labelled SSB-Atto647N (red) bound to ssDNA (**Figure 3G**, top). We assumed that by exclusively exciting DNAp with green light, we would detect SSB fluorescence only when the intermolecular distance falls within the FRET detection range (typically less than ∼10 nm)(*34*, *35*). This methodology enabled us to estimate the relative proximity of the two proteins and probe molecular events in detail. A representative dataset displays the fluorescent trajectories of T7 SSB and T7 DNAp with only the green laser exciting DNAp is shown in **Figure 3H**. Upon correcting for the background noise and signal crosstalk, we identified a transient red SSB signal (**Figure 3H**, top panel). The merged kymograph (**Figure 3H**, bottom panel) further highlights instances of DNAp approaching SSB on ssDNA during replication. Simultaneously, we were able to analyze the replication activity corresponding to these events (**Figure 3G**) using data on DNA elongation (EED). The mechanical measurements revealed that whenever we observe a FRET signal (light blue shaded area), the base pair count remained relatively constant, indicating pausing, or decreases which suggests backtracking. We note that not every pause or backtrack event produced a FRET signal, possibly due to the re-association of unlabelled SSB to ssDNA, and the rapid exchange of labelled DNA polymerase with unlabelled DNA polymerase from the solution(*29*).

**Figure 3I** shows a selection of traces with FRET signals. Mostly, signal was detected during DNAp pausing (75%; N= 6). Notably, we see also FRET during backtracking (25%, N=2). This may indicate the binding of new SSB proteins to the ssDNA strands produced during backtracking. The detected FRET signal during DNAp pausing and backtracking supports our hypothesis of an active SSB displacement model. No FRET signal was observed during the replication process. It should be noted that our FRET signal provides an estimate of relative distance changes rather than absolute distances. The unconventional FRET pairs used in this study (SNAP-Surface® 549 and Atto 647N) might introduce uncertainties in the calculation of absolute distances due to potential variability in the alignment of the donor and acceptor dipoles.

### The C-Terminal Tail and Saturated SSB States: Essential for DNA Replication Dynamics

The C-terminal tail of SSB for effective DNA replication has been reported earlier (*37*, *38*), yet the precise mechanism and the intricate dynamics of SSB interaction with T7 DNAp and its subsequent impact on the replication process remain unclear. Free energy profiles have demonstrated that the local minimum is related to the dissociated state in the DNAp and SSB complex (**Figure 3F**). Here we constructed the free energy profile of the C-terminal truncated variant of T7 gp2.5 (mut T7 SSB) binding to the ssDNA during DNA replication using MD simulation (**Figure 4A**). The blue-colored local energy minima shift from the dissociated state (the position of the right side as indicated in **Figure 3D**) to the bound or intermediate state side (the position of the left side as indicated in **Figure 3D**). This MD simulation revealed that the presence of the C-terminal facilitated the energy transition from bound to dissociated states, suggesting the DNAp might strip SSB of the DNA using the contact with its C-terminal. Moreover, removing the C-terminal significantly weakened the intermediate state due to the C-terminal’s role as an anchor for binding the front basic patch region of DNAp(*33*), which contains four positively charged residues (K587, K589, R590, R591) facilitating strong electrostatic interactions with the C-terminal of SSB. A zoom-in view of the representative structure from the dissociated state in DNAp-SSB energy profile shows the strong contact between the basic patch region and the C-terminal of SSB. The distance between R590 in DNAp and E214 in SSB is 5.6Å, suggesting that strong electrostatic interaction forms. The flanks of the basic patch region, K594 and R599 may also help the binding process of SSB, since R599 also contributes to the electrostatics effects with SSB E222 (**Figure 4B**). Unbiased simulations comparing full-length gp2.5 and the Δ21C mutant with ssDNA highlighted the detachment dynamics, with the mutant displaying increased ssDNA affinity over time (**Figure 4C**). The trajectory movies of the unbiased DNAp-SSB runs (**Movie S2**) and the DNAp-SSBΔ21C runs (**Movie S3**) provides insight into how DNAp actively strips SSB on the DNA replication.

**Figure 4.**
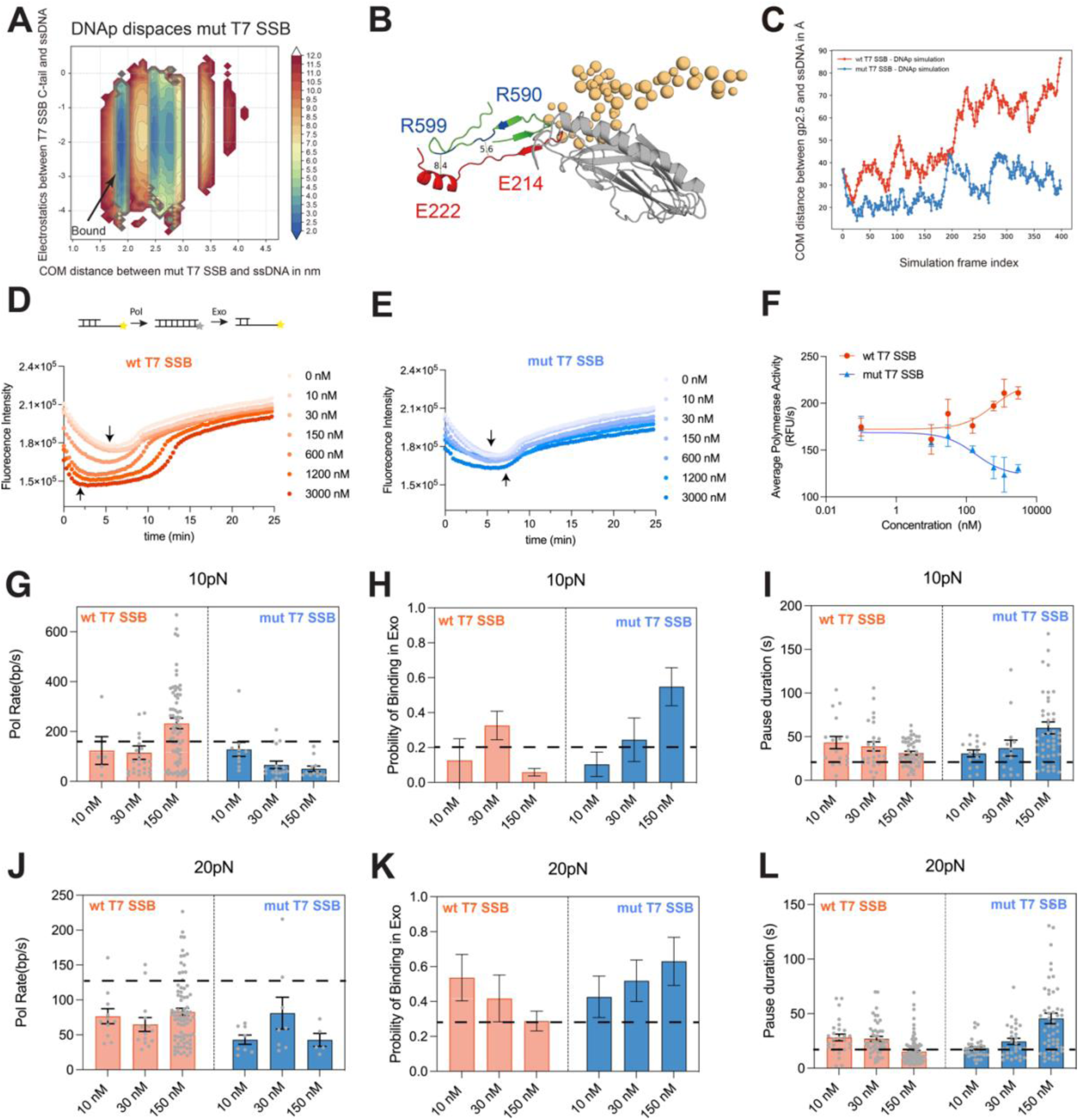
Essentiality of SSB Ratio and C-Terminal Tail for Functional Interactions in Replication Dynamics. **(A)** The two-dimensional free energy profile of the T7 SSB-Δ21C binding to the ssDNA with the existence of DNAp and trx. The barrier among different states has increased from ∼3 kcal/mol to ∼10 kcal/mol and makes it hard to transit among states revealed in Figure 3F. **(B)** A zoom-in view of the C-terminal of SSB binding to the DNAp during the simulation. Orange color shows parts of the coarse-grained ssDNA. Red color shows the C-terminal of SSB. Green color shows partial structures of the front basic patch region of DNAp. The key positive residues in this region (K587, K589, R590, R591, K594, and R599) are colored in blue. The distances between R590-E214 (5.6 Å) and R599-E232 (8.4 Å) are shown. **(C)** The distance between DNA and SSB along the unbias simulations on DNAp+gp2.5 and DNAp+gp2.5Δ21C. The blue color shows the result from DNAp+gp2.5 simulation while orange color shows the result from DNAp+gp2.5Δ21C simulation. Distance variation between DNA and SSB in unbiased simulations of DNAp+gp2.5 and DNAp+gp2.5Δ21C, showing detachment of full-length gp2.5 from ssDNA over time, unlike the gp2.5-Δ21C mutant. **(D)** Real-time DNA primer extension assay monitoring the activity of T7 DNA polymerase with escalating concentrations of wild-type SSB. The wild-type SSB concentrations were 0 nM, 10 nM, 30 nM, 150 nM, 600 nM, 1200 nM, and 3000 nM, with 70 nM DNA and 30 nM DNA polymerase. Black arrows indicate the minimum fluorescence intensity, indicating the transition from pol-dominated to exo-dominated process (N = 3 replicates for each concentration). **(E)** Assay as in **(D)** but utilizing mut T7 SSB across the same concentration gradient (N = 3 replicates for each concentration). **(F)** Quantification of T7 DNA polymerase activity as a function of varying SSB concentrations. The activity was estimated from the initial linear section of the curve (approximately the first 1.5 minutes), representing the polymerase-dominated phase where active DNA synthesis occurs. **(G, J)** Analysis of replication rates (bp/s) for T7 SSB and mut T7 SSB across various concentrations under forces of 10pN (H) and 20pN (K), respectively. Experiments utilized 30 nM DNA polymerase and varying concentrations of wild-type or mutant T7 SSB (10, 30, 150 nM), quantified via single-molecule force spectroscopy with replication rate determined by a change-point detection algorithm analyzing base-pair traces from a dual-optical trapping system. Horizontal dashed line shows the replication rate without SSB as a reference (data from Figure 1D). Sample sizes: under 10 pN force, N = 5 (10 nM), N = 25 (30 nM), N = 72 (150 nM) for T7 SSB; N = 10 (10 nM), N = 17 (30 nM), N = 12 (150 nM) for mut T7 SSB; under 20 pN force, N = 11 (10 nM), N = 14 (30 nM), N = 81 (150 nM) for T7 SSB; N = 8 (10 nM), N = 8 (30 nM), N = 5 (150 nM) for mut T7 SSB. Data presented as mean ± SEM. **(H, K)** Examine the relative probability of DNA polymerase binding at the exonuclease active site with differing T7 SSB and mut T7 SSB concentrations at 10pN **(I)** and 20pN **(L)** forces, respectively. Probability calculated by the ratio of exonuclease events to total exonuclease and polymerase events per DNA molecule. Horizontal dashed line shows the replication rate without SSB as a reference (data from Figure 1E). Sample sizes: under 10 pN force, N = 4 (10 nM), N = 11 (30 nM), N = 47 (150 nM) for T7 SSB; N = 8 (10 nM), N = 9 (30 nM), N = 12 (150 nM) for mut T7 SSB; under 20 pN force, N = 10 (10 nM), N = 8 (30 nM), N = 43 (150 nM) for T7 SSB; N = 9 (10 nM), N = 9 (30 nM), N = 12 (150 nM) for mut T7 SSB. Data presented as mean ± SEM. **(I, L)** Compare average pause durations of DNA replication with differing T7 SSB and mut T7 SSB concentrations at 10pN (J) and 20pN (M) forces, respectively. Horizontal dashed line shows the replication rate without SSB as a reference (data from Figure 1G). Sample sizes: under 10 pN force, N = 10 (10 nM), N = 26 (30 nM), N = 50 (150 nM) for T7 SSB; N = 15 (10 nM), N = 14 (30 nM), N = 47 (150 nM) for mut T7 SSB; under 20 pN force, N = 27 (10 nM), N = 48 (30 nM), N = 99 (150 nM) for T7 SSB; N = 32 (10 nM), N = 30 (30 nM), N = 52 (150 nM) for mut T7 SSB. Data presented as mean ± SEM.

Further, we examined the C-terminal’s role in modulating polymerase activity across varying SSB-DNA saturation levels using real-time primer extension assays(*39*). By employing real-time primer extension assays enabled by a 5′-end fluorophore, quenched upon nucleotide incorporation, we monitored the kinetics of DNA synthesis in real-time (**Figure S8A-B**). Wild-type SSB addition accelerated the polymerization rate, as evidenced by a shortened polymerization phase with increasing SSB concentrations (black arrow shift left with increasing concentration, **Figure 4D**), suggesting a shorter time to complete the DNA synthesis. In contrast, the mutant SSB showed a slightly reduced polymerization rate (**Figure 4E**). We then compared the relative polymerase activities, varied with SSB concentration, by reporting initial rates (RFU/s) as a reasonable approximation measured during the polymerase-dominated phase (**Methods**, **Figure 4F**). These results demonstrate that the presence of wt T7 SSB increases the replication rate (increased by ∼20% from 0nM to 3000nM wt T7 SSB), while the presence of mutant SSB leads to a decreased rate (decrease by ∼25% from 0 nM to 3000 nM mut T7 SSB). Notably, this effect becomes apparent only at concentrations higher than 150 nM (**Figure 4F**). Given the footprint of wt T7 SSB of ∼ 10 nt per SSB molecule(*20*), our experimental results indicate that saturation occurs near a 2:1 ratio of SSB to DNA, which aligns with the requirement of 150nM SSB to achieve saturation for 70nM ssDNA with a 17 nt overhang (**Methods**). This result suggests that SSB may “pull” DNAp forward during replication especially under saturated SSB conditions, while the SSB mutant seems to hinder replication. This confirms that the SSB’s C-terminal tail plays a crucial role in facilitating DNAp’s progression along the ssDNA template.

T7 DNA polymerase possesses an intrinsic exonuclease domain(*36*), facilitating proofreading. Correspondingly, our real-time assay trace demonstrated a subsequent increase in fluorescence intensity, indicative of exonuclease activity (**Figure 4D-E**). As a control, we observed that the fluorescence signal remains relatively stable over the experimental timeframe (∼30 min) when only SSB is added to DNA. However, higher SSB concentrations correlate with decreased overall fluorescence (**Figure S8C-E**). This effect might be attributed to SSB binding to ssDNA, causing fluorescence signal quenching. Importantly, this control experiment enables reliable quantification of SSB effects on DNA polymerase activity.

To further elucidate the dynamic mechanism of DNAp interacting with the C-terminal truncated SSB under physiologically relevant tensions, we employed force spectroscopy to measure and detect the interaction between DNAp and mut T7 SSB at varying concentrations and compared the dynamics with that of the T7 SSB. Our previous findings(*20*) have indicated that T7 SSB reaches saturation on DNA binding, in single-DNA molecule assays, at concentrations exceeding 50nM. We examined the replication and exonuclease activities of DNAp under three distinct SSB concentration regimes: low (10nM), semi-saturated (30nM), and fully saturated (150nM), utilizing single-molecule, high-resolution optical tweezers to track DNA length alterations.

At a force of 10 pN, the replication rate was enhanced in the presence of fully saturated wt T7 SSB compared to DNAp replication without SSB (230 ± 20 bp/s v.s.180 ± 10 bp/s, horizontal dashed line from **Figure 1D** added as a reference, **Figure 4G**, mean ± SEM). This contrast was not clearly observed at low or semi-saturated SSB levels (120 ± 50bp/s, and 120 ± 30 bp/s, respectively, mean ± SEM). Conversely, the presence of mutant SSB reduced the polymerase rate in a concentration-dependent manner, corroborating prior real-time DNA primer extension analyses (**Figure 4F**). We quantified the likelihood of polymerase engaging in exonuclease activity (**Figure 4H**), revealing that wt T7 SSB, particularly at saturated levels, preferentially positions the polymerase in polymerase state (0.1 ± 0.1 at 10nM vs. 0.06 ± 0.02 at 150nM, mean ± SEM). In contrast, the C-terminal tail-lacking mutant SSB favors exonuclease state binding (0.1 ± 0.06 at 10nM v.s. 0.6 ± 0.1 at 150nM, mean ± SEM). This effect can be attributed to two factors: the C-terminal domain’s interaction with DNAp, possibly through the trx domain from DNAp, facilitating DNAps’ forward movement and reducing proofreading occurrences, and wt T7 SSBs’ reduced DNA affinity, minimizing replication hindrance. The T7 mut SSB on the other hand binds to ssDNA more firmly but fails to interact constructively with DNAp, i.e. it seems mostly to act as a roadblock. This suggests DNAp might be only able to advance when mutant SSB disassociates, as further evidenced by increased pausing times in mutant SSB (**Figure 4I**) (30 ± 3s at 10nM v.s. 60 ± 6s at 150nM, mean ± SEM). The presence of wt T7 SSB also increases pausing times of DNA polymerase (**Figure 4I**, reference dashed line from **Figure 1G**). This supports the hypothesis that SSB bound to ssDNA recruits DNA polymerase to the DNA template, creating a roadblock that causes the replicative polymerase to pause (**Figure 1H**). Interestingly, lower concentrations of SSB lead to longer pauses (40 ± 10 s at 10nM) compared to higher concentrations, which result in shorter pauses (30 ± 2 s at 150nM), as shown in **Figure 4I**. This observation might be due to the fact that at lower SSB concentrations, the protein is located at random positions on the ssDNA, which can randomly recruit DNA polymerase to bind to the ssDNA and cause longer pauses. In contrast, at higher, saturating SSB concentrations, there is less available space for the polymerase to bind, leading to shorter pausing durations.

Under conditions promoting less secondary structure (20pN), both the presence of wt T7 SSB and mut T7 SSB decreased replication rates (**Figure 4J**), consistent with earlier findings in **Figure 1**. Assessing exonuclease site binding probabilities revealed that mutant SSB presence increases the likelihood of exonuclease state binding with rising SSB concentrations (**Figure 4K**) (0.4 ± 0.1 at 10nM vs. 0.6 ± 0.1 at 150nM). Interestingly, saturated wt T7 SSB conditions showed similar exonuclease binding levels as in the complete absence of SSB. While a low concentration of SSB increased the probability of binding in the exo-state (0.5 ± 0.1 at 10nM vs. 0.3 ± 0.05 at 150nM). These results indicate that at increased tension, any SSB presence acts as a blockade, leading to lower replication rates and a higher probability of DNA polymerase binding in the exonuclease state. This is likely due to the reduced need to compete with the secondary structure of ssDNA. As expected, at lower wt T7 SSB concentrations, the protein is more likely to recruit and stabilize DNA polymerase on the ssDNA, forming a self-imposed roadblock that further reduces the replication rate and increases the exonuclease state probability. This is further supported by the analysis of average pausing durations (**Figure 4L**), where lower wt T7 SSB concentrations (10 nM) resulted in longer pauses (28 ± 3 s) compared to no SSB (13 ± 1 s), while saturated wt T7 SSB levels (150 nM) produced pause times comparable to those observed without SSB (15 ± 1 s). Interestingly, the mutant SSB binds more firmly to ssDNA but fails to interact constructively with DNA polymerase, primarily acting as a roadblock. Higher concentrations of the mutant SSB thus increased the DNA polymerase blockage times (18 ± 1 s at 10 nM vs. 45 ± 5 s at 150 nM), demonstrating a concentration-dependent effect.

## Discussion

Genomic DNA is a crowded molecular environment where DNA motor proteins frequently collide. A notable example is the encounter between replicative DNAp and high-affinity SSBs coating ssDNA during DNA replication. In this study, we investigated the molecular mechanisms and visualized the dynamic process by which DNAp overcomes these SSBs, resolving molecular collisions on DNA and elucidating the molecular consequences of this interaction. Our results support a sequential and active displacement model, where SSBs are displaced one after another by the advancing DNAp through functional domain interaction. This coordinated process not only enables efficient SSB removal but also promoting each SSB encounter to enhance overall replication efficiency, suggesting an evolved synergy between these seemingly antagonistic proteins.

Our single-molecule dual-color imaging experiments provide compelling evidence for this mechanism. Measurements of the diffusion constant of T7 SSB on ssDNA confirm their relative stability on the template, consistent with previous findings(*13*), suggesting that SSBs are not easily displaced in clusters. High occupancy of SSBs on the template restricts individual SSB diffusion, and the lack of free ssDNA ends in Okazaki fragments prevents SSBs from being pushed off by the advancing lagging-strand polymerase. These observations together support the sequential displacement model during lagging-strand DNA synthesis. This sequential displacement contrasts with alternative mechanisms proposed in other systems(*17–19*, *40*), where SSBs might be removed or bypassed through different processes. Such mechanisms may involve helicases XPD displacing SSBs(*17*), alterations in SSB binding modes(*40*), or translocases advancing along ssDNA by pushing SSBs forward(*18*). These variations suggest that SSB proteins from different species—with distinct diffusion properties, binding modes, physiological functions, and biological contexts—may interact differently with DNA translocases. In our system, sequential displacement ensures that SSBs remain bound to ssDNA until DNAp arrives, providing continuous protection against nucleases and preventing secondary structure formation(*1*, *2*), thereby minimizing the risk of DNA damage or replication errors.

Using AWSEM coarse-grained MD simulations, we modeled the DNAp-SSB interaction, revealing strong electrostatic interactions between the basic patch on the front of DNAp and the acidic C-terminal tail of SSB. Our experimental findings, using single-molecule FRET and high-resolution force measurements, suggest direct contact and regulatory mechanisms between these proteins. The observed FRET signals imply direct interactions facilitating the displacement of SSB by DNAp during replication. Notably, this displacement does not perturb the overall structure of DNAp’s interaction with thioredoxin (trx), in line with conclusions from prior research(*41*). The highly negative charge of SSB’s C-terminus, akin to the PEST sequence in the C-terminus of IκBα, suggests a universal mechanism among DNA regulatory proteins(*42*, *43*), involving molecular stripping facilitated by electrostatic interaction. This interaction possibly alters the binding affinity of SSB, leading to its dissociation and facilitating the forward movement of DNAp. Disruptions to this interplay, as observed with tailless SSB, result in diminished DNA replication rates.

Our single-molecule investigations reveal that SSB proteins play a dual role in DNA replication, dependent on the ssDNA conformation regulated by biological relevant forces. Under low-tension conditions (forces < ∼ 15 pN), where ssDNA is prone to forming secondary structures, SSBs prevent these formations, smoothing the DNA template for replication and increasing processivity, aligning with previous studies(*4*, *10*). Furthermore, the electrostatic interaction between the acidic C-terminal tail of SSB and the basic patch on DNAp increases the replication rate, potentially by facilitating the forward movement of DNAp. The increased probability of DNAp binding in the polymerase-active direction in the presence of SSB indicates that SSBs promote DNAp’s active conformation, likely through direct physical interactions. These single-molecule observations are consistent with prior in vitro ensemble studies conducted under low- or no-tension conditions, which have demonstrated a notable acceleration in DNA replication in the presence of SSBs(*3*, *9*, *10*, *44–46*). The observed increase in replication rate with wild-type T7 SSB, particularly at saturating concentrations, suggests that each individual SSB molecule assists in facilitating DNAp progression along ssDNA. This facilitation is likely mediated by the C-terminal tail of SSB, as mutants lacking this domain did not enhance the replication rate(*37*, *38*). According to the proposed active displacement mechanism, DNAp advancement coordinates with SSB dissociation by lowering the energy barrier, effectively regulating SSB removal during replication.

Conversely, at higher tensions (> ∼20 pN), where secondary structure formation is less likely, SSBs may act counterproductively as physical barriers, reducing replication efficiency. One possible explanation is that SSBs function as weak blocks under both low and high forces; however, under low forces, the more substantial roadblocks formed by hairpin structures overshadow the obstruction created by SSBs. Therefore, SSBs promote replication under low force by preventing the formation of larger secondary structures that would impede DNAp. At high forces, where hairpin formation is suppressed, the obstruction from SSBs becomes more significant. Additionally, excessive recruitment of DNAp due to SSBs positioned along ssDNA can create self-imposed obstacles, leading to extended pause durations during replication. This suggests that while SSBs are beneficial under certain conditions, they may inadvertently impede replication by acting as hindrances under others. The force-dependent impact of SSBs implies a regulatory mechanism mediated by biological tension—possibly induced by other molecular motors or DNA constraints(*47*)—allowing cells to modulate replication dynamics in response to cellular conditions or stress. Our findings align with similar observations in *E. coli*(*6*) and mitochondrial(*4*) systems, affirming the integral role of SSBs in modulating replication efficiency.

Positioning our findings within the broader biological context highlights the critical role of SSBs in DNA replication dynamics. Yet, various potential discrepancies exist between our experimental conditions and the in vivo situation. Our research focused on an in vitro phage system, which may oversimplify cellular replication complexities. *In vivo*, replication involves intricate systems, potentially including hybrid machinery from both viral and host cell replisomes. Due to limited temporal resolution in confocal fluorescence microscopy, we did not address how rapid DNAp exchange is affected by SSB displacement. The low throughput of single-molecule assays also limited our survey of other SSBs. Future studies should extend investigations to other replicative systems, such as eukaryotes with distinctive SSB diffusive properties. Such studies would more closely mimic *in vivo* conditions, broadening the applicability of our findings and potentially uncovering universal principles of DNA replication regulation.

## Materials and Methods

### DNA Template Construction, Protein Purification and Labelling

The pKYB1 vector (∼8.3 kb) DNA construct with biotin labels at both ends was generated using established protocols(*48*). Recombinant T7 DNA polymerase (T7 gp5 fused with trxA) was expressed, purified, and labeled with SNAP-Surface® 549. Pure protein and labeled protein were aliquoted, flash-frozen, and stored at -80°C. Wild type T7 gp2.5 were purified and labeled with Atto647N using a protocol based on previous literature(*49*).

### Single Molecule Set-up

Single-molecule experiments were performed on a LUMICKS C-Trap instrument, which combines three-color confocal fluorescence microscopy and dual-trap optical tweezers with a 5-channel microfluidic flow-cell. A 1064 nm fiber laser was used via a water-immersion microscope objective to generate two orthogonally polarized optical traps. The biotinylated pKYB1 DNA construct was tethered between two 1.76 μm streptavidin-coated microspheres (Spherotech Inc) in situ within the flow cell. A single DNA molecule was detected and confirmed through a change in the F-x curve. Orthogonal channels 4 and 5 were used as protein loading and/or experimental imaging chambers as described for each assay. All experiments were performed at room temperature with force data collected at 50 kHz. Fluorescently labeled DNA polymerase was excited at 532nm and fluorescently labelled T7 SSB was excited at 638nm, and to optimize imaging data over a long measurement period, the protein was scanned with a pixel dwell time of 1-2 ms, laser power of 1.2mW (30%), and scanning interval of 8ms unless stated otherwise.

### Experimental Assay

T7 DNA polymerase concentration was set at 30 nM with a labelling efficiency of ∼60%. T7 gp2.5 concentration was set at 150nM with a labelling efficiency of ∼4%. The fluorescently labelled SSBs, mixed with unlabelled SSBs, served as markers to determine the relative distance between the replicative DNA polymerase and the SSB proteins.

To prevent nonspecific binding, the flow cell was cleaned with bleach and passivated overnight with 0.1% BSA and 0.5% Pluronic. The experiments were conducted in a standard measurement buffer composed of 20mM Tris-HCl pH 7.5, 100mM NaCl, 5mM MgCl2, 1mM DTT, and 0.02% BSA.

Single DNA molecules were captured and moved into a protein channel containing DNA polymerase and SSB. The activity of the DNA polymerase was studied by monitoring the change in DNA extension at a constant tension of 40-50pN. This tension was used to digest ∼ 5 kbp of dsDNA and create a long section of ssDNA as a template for polymerization activity. Tension was then decreased to 10-20pN to measure polymerization activity in the presence or absence of SSB.

### Coarse-grained Force Field for Protein-ssDNA System

The latest iteration of the protein folding force field, known as the Associative memory, water-mediated, structure and energy model (AWSEM), builds upon a sequence of models refined through energy landscape theory and a quantitative machine learning approach for protein folding. AWSEM represents each amino acid residue using three actual atoms (CA, CB, and O) and three virtual sites (C, N, H, with exceptions for proline and glycine). The AWSEM-Suite, an updated version of AWSEM, has demonstrated its predictive accuracy in the CASP13 protein structure prediction competition, achieving top three server predictions in several instances(*50*).

In our previous work of helicase gp4-ssDNA system, we have extended AWSEM to a transferrable protein-DNA model with a modified 3SPN.2C force field that have shown the physical properties of ssDNA(*24*, *51*). Our coarse-grained DNA model incorporates detailed features of DNA structure, including specific hydrogen bonding and base stacking interactions characteristic of the B-DNA form. Similar to AWSEM, the DNA force field uses three beads to indicate the phosphate, sulphur, and nitrogen base atom group.

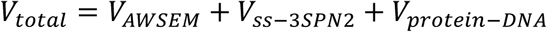

For the details of the AWSEM and ss-3SPN2 energy composition, please refers to Ref25. Briefly, AWSEM includes.

### Construct the Initial Structures for MD Simulation

The currently only available gp2.5 structure in PDB is 1JE5, which lacks the most flexible C-terminal region (residue 203 - 232) and the linker region (residue 79 - 88)(*31*). Deep-learning based protein structure prediction model Alphafold2 and Modeller version 9.23 were used to construct the full-length gp2.5 structure(*32*, *52*). AlphaFold2 is a state-of-the-art transformer-based model developed by DeepMind that can predict protein structures from amino acid sequences with atomic-level accuracy. In contrast, Modeller represents the traditional template-based modelling tool for protein structure that achieves good performance. Since the C-terminal is tightly bind to the positive binding groove in the Alphafold2 predicted structure, a short simulation with external force to expel the C-terminal tail from the groove was did to probe an initial state that ssDNA have balanced probability to bind.

For the gp2.5-gp5-ssDNA complex, we choose structure PDB ID 6P7E as a template because this structure contains a peptide DTDF that mimics the C-terminal tail of gp4(*33*). By aligning the C-terminal of the gp2.5 structure to the short peptide and extending the 5’ end of the ssDNA with poly-A, we create an initial complex that gp2.5 closes to the longer ssDNA chain. We also use the inferred structures from SAXS data in the Figure 7 from Foster et al. as a reference(*33*).

### Free Energy Calculations Based on the Umbrella Sampling Technique

Umbrella sampling is an enhanced sampling technique that could force the exploration of regions of state space that would otherwise have insufficient sampling. It uses a series of independent windows along a selected collective variable, which serves as a continuous parameter to describe the system from a higher dimensional space to model a conformational transition(*53*). In the current system, we use the center of mass between gp2.5 and the first 20 nucleotides of the ssDNA as one metric while the electrostatics between the charged residues in the positive binding pocket of gp2.5 (residue 72-94, 113-125, 157-165) and ssDNA as the second metric. For the DNAp-SSB complex, we used the distances between the COM of the SSB and the COM of the first 21 base pairs of the 5’ overhang ssDNA as the biasing coordinates to build free energy profiles. No external bias was added to the interaction of DNAp and T7 SSB.

The harmonic biasing potential used for constant temperature umbrella sampling simulations for 8 million steps was scaled to 10 kcal/mol. The biasing center values were chosen to be equally spaced from 12.5 Å to 27 Å with an increment of 0.5 Å and 50 windows. The initial structures for umbrella sampling were picked as the complex structure described in the above section. The weighted histogram analysis method (WHAM) is used to reconstruct the unbiased free energy landscapes from the umbrella sampling data(*54*). An additional dimension related to the specific conformation changes of interest was added for plotting the 2D free energy profile.

### Protein Remote Homology Detection

The protein remote homology detection is conducted by HHpred (https://toolkit.tuebingen.mpg.de/tools/hhpred)(55). The default structural/domain databases (PDB_mmCIF70_8_Mar) was used to perform the search.

HHpred simplifies sequence database searching and structure prediction, offering a user-friendly experience akin to BLAST or PSI-BLAST while significantly enhancing sensitivity in identifying distant homologs. HHpred scans alignment databases such as Pfam or SMART, streamlining search results into sequence families rather than individual sequences. The method operates on the comparison of profile hidden Markov models (HMMs), contrasting with traditional sequence search methods that scan databases like UniProt or NR.

### Structure Clustering and Heatmap Analysis

The structure frames that fall within the local energy basin area during the umbrella sampling were extracted based on the free energy profile. The Qw value was used to measure the pairwise distance between two structures, with a detailed explanation provided in(*50*). After calculating the Qw value for all structures in the local basin, the hierarchically clustered heatmap and dendrogram were computed using the clustermap function in the seaborn package and the dendrogram function in the scipy package, respectively. The Y axis indicates the distance of each branch, as computed by the scipy.cluster.hierarchy.linkage package. The center of the largest cluster was then determined and manually picked out for further analysis.

### Real-time DNA primer extension assay

The activity of T7 DNA polymerase was measured in the presence or absence of wild-type T7 single-stranded DNA-binding protein (T7 gp2.5, wt T7 SSB) and a deletion mutant lacking 21 C-terminal residues (T7 gp2.5-Δ21C, mut T7 SSB) using a 45-nt long DNA template annealed to a 28-nt primer strand. The carboxyfluorescein (6-FAM) moiety was located at the 5′ end of the template strand. Template strand: 5′/6-FAM/-CCCCCCCCCATGCATGCGCACCTAAAGTTGGGAGTCCTTCGTCCTA-3′. Primer strand: 5′-TAGGACGAAGGACTCCCAACTTTAGGTG-3′. Reactions were performed in a final volume of 10 μl with 25 μm of each dNTP in buffer 20 mM Tris–HCl pH 7.5, 50 mM NaCl, 3mM MgCl2, 1 mM DTT, 0.05% BSA and 0.05% Tween-20. Reactions were initiated by the addition of 70 nM labelled DNA to 30 nM DNA polymerase pre-mixed with indicated concentrtion of wt T7 SSB or mut T7 SSB and measured in a 348-well plate using a BMG Labtech Pherastar FS plate reader during 30 min at 25°C.

The polymerase activity was estimated from the initial linear section of the curve (approximately the first 1.5 min), representing the polymerase-dominated phase where DNA synthesis actively occurs. The slope derived from this fit gives an estimate of the initial polymerase activity in RFU/s. Data analysis was performed using self-written python script and GraphPad Prism software.

### Single Molecule Experiment Data analysis

All data obtained from C-trap is in. tdms format and is analyzed using custom-written Python scripts. Details for access is provided in **CODE AVAILABILITY** section.

### Analysis of ssDNA/dsDNA Junction Trajectory

In our investigation, we manipulated the template tension using an optical tweezer system. The T7 DNA polymerase was tailored to function as an exonuclease at tensions ranging from 40-50pN, producing partially ssDNA, and to execute polymerization at tensions between 10-20pN, facilitating the synthesis of dsDNA. Owing to the catalytic activity of DNA polymerase, the end-to-end distance (EED), measured directly between two optically entrapped beads, comprising both ssDNA and dsDNA, was consistently monitored over time.

We applied a composite of the Freely Jointed Chain (FJC) and Worm-Like Chain (WLC) models to determine the proportions of ssDNA and dsDNA present within the EED, respectively. Given the elasticity differences between ssDNA (described by an FJC model(*56*)) and dsDNA (described by an extensible WLC model(*57*)), the ssDNA percentage can be derived using the equation:

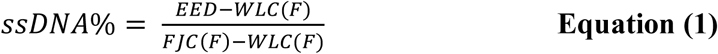

To adjust for the changing ssDNA length during DNA replication involving saturated SSB binding, we employ a correction factor. This factor accommodates the transient shortening of ssDNA caused by SSB binding during DNA polymerase activity.

The correction formula applied is:

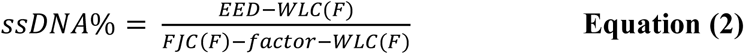

Here, ‘EED’ represents end-to-end distaces between the two optically trapped beads, while ‘FJC’ and ‘WLC’ denote the Freely Jointed Chain(*56*) and Worm-Like Chain(*57*) models, respectively. ‘F’ corresponds to the applied force. The correction factor accounts for the ssDNA length alteration when fully coated by SSB proteins, estimated as 0.246 µm under 10pN and 0.177 µm under 20pN with gp2.5 protein (**Figure S2**). This methodology refines our ssDNA length estimates during replication, enhancing the precision of DNA replication dynamics analysis.

The dsDNA percentage can be calculated as

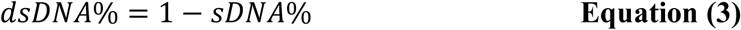

The ssDNA/dsDNA junction position can be determined by assuming that ssDNA appears on the top half of the kymograph:

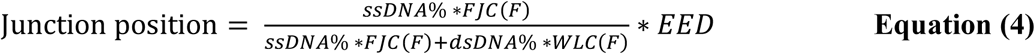

If the dsDNA appears on the top half of the kymograph, the junction position can be derived as:

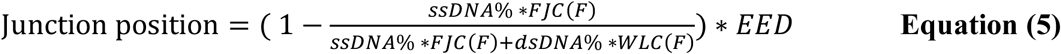

This method facilitates high-resolution, real-time tracking of DNAp’s position, even amidst sporadic dissociation of the fluorescently labeled DNAp.

### Analysis of Single-Molecule Basepair-Time Traces

We employed a pwlf Python library(*58*) to fit piecewise linear functions to single-molecule base pair-time traces and detect the most likely change points, which signify shifts in polymerase (pol) or exonuclease (exo) activities. To reduce noise, the traces were initially processed with a Savitzky-Golay (SG) filter. The piecewise linear function, adapted from previous publications(*58*) with minor modifications, was then used to detect these change points, allowing us to mark the steps or change points in the base pair-time traces. Subsequently, we quantified the processivity, velocity, and duration of each identified segment.

One crucial aspect of our study involves the detection and characterization of pausing events, denoted by temporary halts in polymerase activity that yield steady basepair values over a significant period. The ‘noise threshold’ is determined as the standard deviation of basepair-time traces in the absence of protein under the tensions under investigation, estimated by employing the median absolute deviation (MAD) of sequential data points and amplified by a constant factor (1.4826). This method facilitates a robust distinction between inherent fluctuations (noise) and genuine pausing events (signal). The ‘significant period’ is deduced from the known exonuclease rate of T7 DNAp (∼100bp/s) and replication rate (∼200bp/s), indicative of the standard duration necessary for the polymerase to add a base pair under our experimental parameters. We define a pause as an event where the actual process duration is 10 times greater than the expected time, which, for simplicity, is defined to occur at 10bp/s under 50pN and 20bp/s under 20pN or 10pN.

### Analysis of Fluorescent Protein Movement using Kymographs

The acquired kymographs were analysed with a custom-written tracking program in Python using lumicks.pylake package. In brief, the line identification algorithm(*59*) was employed to identify the centers of lines by leveraging local geometric principles, while the greedy algorithm was used for initial feature point detection and line tracing(*60*, *61*).

### Analysis of SSB Diffusion using Kymographs

The diffusion constant of T7 SSB was calculated using the Mean Square Displacement (MSD) method, exploiting the extracted fluorescence trajectory, as illustrated by the yellow line in **Figure 2D**. The computed diffusion constant of 7 ± 7 * 10^-4^ μm^2^/s (mean ± SEM) (**Figure 2H**), in agreement with our recent report(*13*), substantiates that the SSB is essentially immobile on ssDNA. By comparison, the known static protein EcoRV was previously determined in the same lab(*30*) to exhibit an apparent diffusion constant of 1*10^-4^ µm^2^/s with a localization accuracy value (∼10 nm, employing integration times of 1 s), slightly lower than that determined in our experiments. This supports the hypothesis that T7 SSB binds at a fixed position and does not diffuse.

### Correlation Between ssDNA/dsDNA Junction with Fluorescently Labelled DNA Polymerase

Alongside a distance-time curve, we recorded the fluorescence trajectory from labelled proteins, enabling real-time tracking of DNA polymerase activity at the ssDNA/dsDNA junction. This was achieved by overlaying the calculated ssDNA/dsDNA junction movement, based on earlier calculation, with the DNAp trajectory in the kymograph. We adjusted for variations in starting time and position between the optical tweezer and fluorescence microscopy systems. Fluorescence intensity was extracted along the junction over time, focusing on the green channel to reduce signal crosstalk. A 5-pixel-wide box was used to capture photons within the point-spread function, ensuring optimal signal collection. An optimal overlap between the calculated ssDNA/dsDNA junction and the fluorescent trajectory was determined by seeking the maximum local intensity. We recalibrated the x-offset and y-offset based on these findings.

### Determination of the Distance Between DNAp and SSB

The optimized DNAp trace was overlaid with the fluorescently labelled SSB. To enable further quantification, the SSB trace was isolated and subjected to trajectory analysis, considering only those SSB trajectories which spanned more than 3 pixels (∼ 1s). These fluorescently captured trajectories were subsequently aligned with the DNAp trajectories, calculated from the ssDNA/dsDNA junction. Following this alignment, the distance between the DNAp and the SSB, was measured. In addition to these distance measures, we also computed the duration, rate, and diffusion constant of SSB to further illuminate the underlying mechanisms of the DNAp-SSB interaction.

### Error Discussions on Tracking the *Real-time* Interaction between DNAp and SSB

Potential sources of error were carefully considered and mitigated in tracking the interaction between DNAp and SSB, which involved tracking the DNAp position, detecting SSB trajectories, and overlapping the optimized DNAp and SSB traces. First, although optimal overlapping of the calculated ssDNA/dsDNA junction and the fluorescent trajectory was attempted, the DNAp track could occasionally deviate from the true trace due to local fluctuations in the fluorescence signal or instrumental noise. Errors may also arise from inaccuracies in estimating the bead size (**Figure S2B**) Calibration procedures were implemented to minimize these errors. Second, in detecting SSB trajectories, systematic errors arising from the point spread function and diffraction limit were addressed by employing a line identification algorithm and a greedy algorithm for initial feature point detection and line tracing (**Methods**). Background subtraction and signal-to-noise ratio optimization techniques were also applied to enhance the accuracy of SSB trajectory detection. Third, in overlapping the optimized DNAp and SSB traces, slight misalignments could occur due to the use of different dyes and excitation lasers (532 nm and 638 nm, respectively) for labeling DNAp and SSB, leading to variations in optical properties and potential crosstalk between fluorescence channels. Appropriate measures, such as spectral unmixing and channel compensation, were taken to minimize these effects. By acknowledging and addressing these potential sources of error, we aimed to minimize their impact, thereby enhancing the reliability and reproducibility of our findings on the determination of the distance between DNAp and SSB.

### Quantification and Statistical Analysis

The number of molecules or events analyzed is indicated in the text or figure legends. Errors reported in this study represent the standard error of the mean, unless otherwise stated. *p* values were determined from unpaired two-tailed t-tests using GraphPad Prism 9 (ns, not significant; *p* values are indicated in the figure legends). All experiments were independently repeated at least three times with similar results. Representative results are shown in figures.

## Acknowledgments

We thank Seyda Aca and Sandrine D’Haene for assistance with protein purification and DNA construction, Noémie Danné for help with implementing the step-fitting algorithm. We thank C.C Richardson for providing us the gp2.5-Δ21C samples from his lab. We thank Chen-Yu Lo and Yang Gao from Rice University for discussion. We thank Erwin Peterman for critical reading and constructive feedback of this manuscript.

## Author contributions

L.X, S.J. and G.J.L.W. conceptualized the research. L.X prepared protein samples, collected and analyzed the single-molecule data; S.J. implemented the MD simulation and analyzed the data; L.X and M.U. collected and analyzed the real-time DNA primer extension data. S-J L. provided purified gp2.5-Δ21C and tested their biochemical activity; L. X., S.J. and X.C. wrote the original manuscript; L. X., S.J., P.G.W., and G.J.L.W. edited the manuscript; G.J.L.W. and P.G.W supervised the project; the manuscript is read and confirmed by all the listed authors.

## Data and materials availability

The datasets generated and/or analyzed during the current study will be uploaded to public repository upon conditionally accepted.

The custom-written python scripts for analyzing basepair-time traces, for analyzing the displacement of SSB and for analyzing the real-time DNA primer extension data are available from Github: https://github.com/longfuxu/Interplay_Between_DNAPol_and_SSB, under the MPL-2.0 license. The script for analzying the MD simulation result and the Weighted Histogram Analysis Method (WHAM) are available from GitHub: https://github.com/CryoSky/MD_simulation_DNAp_SSB. The repository includes example dataset, example Jupiter notebook, along with a detailed README file for instructions on installation and usage.

**Figure S1.**
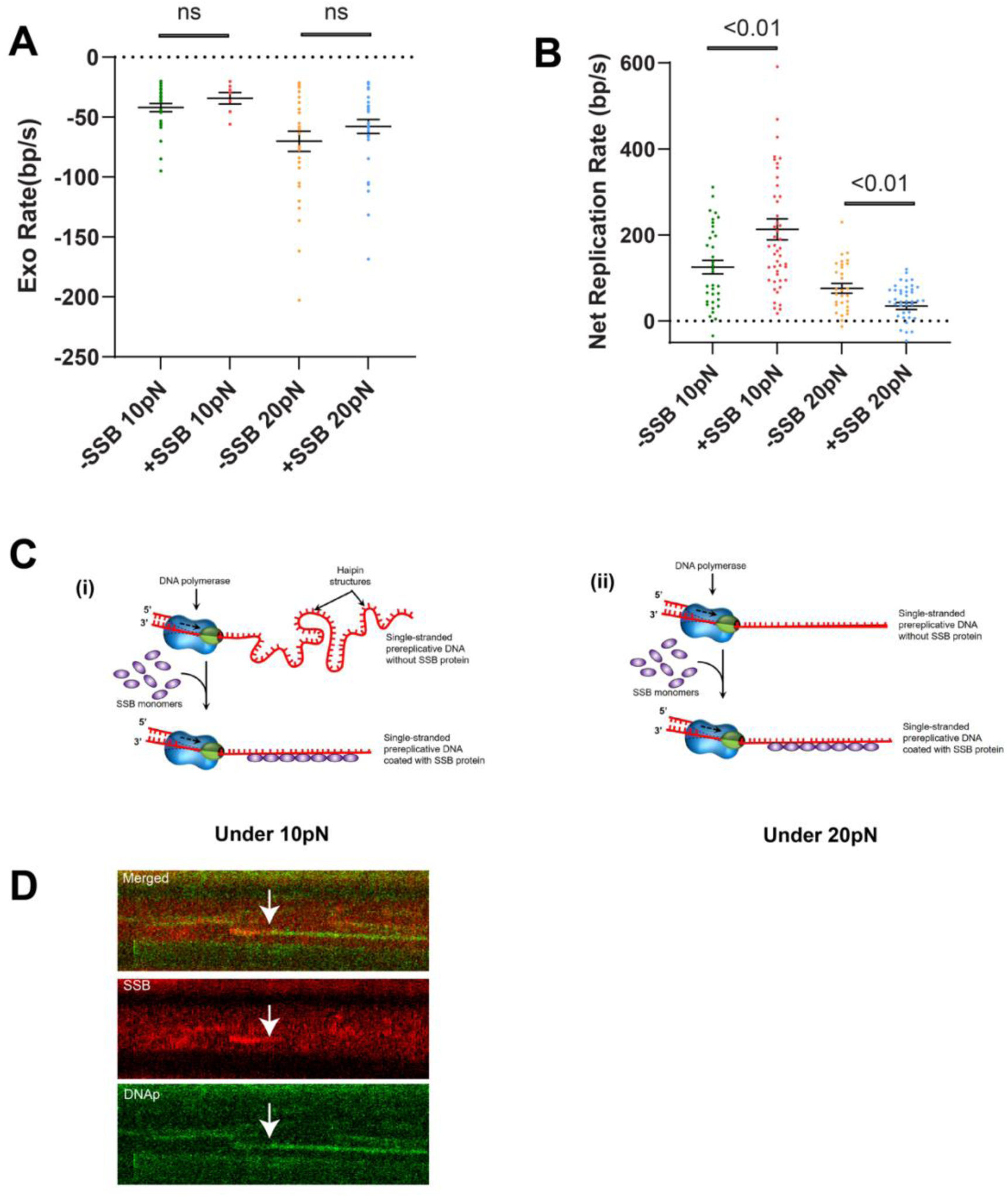
SSB’s effect on proofreading activity and net replication rate. **(A)** Illustration of SSB’s effect on polymerase’ proofreading activity (exo rate) under varying DNA template tensions. Exo rate slightly increases from -42 ± 3 bp/s (N=29) to -34 ± 5 bp/s (N=7) at 10 pN with the presence of SSB and increases from -70 ± 8 bp/s (N=31) to -57 ± 6 bp/s (N=33) at 20 pN. Data are collected from 30-47 different DNA molecule cycles; data is represented as mean ± SEM. **(B)** Depicting net replication rate with or without SSB at 10 pN and 20 pN. In the presence of SSB, net replication rate increases from 125 ± 15 bp/s (N=35) to 213 ± 24 bp/s (N=47) at 10 pN, while decreases from 76 ± 11 s (N=30) to 34 ± 8bp/s (N=43) at 20pN. Data are collected from 30-47 different DNA molecule cycles; data is represented as mean ± SEM. **(C)** Schematics showing the role of SSB proteins in preventing secondary structures in intermediate ssDNA during replication and facilitating DNA polymerization. (i) At 10pN force, where secondary structures can form, DNAp accelerates in the presence of SSB; (ii) as a comparison, replication decelerates under similar conditions at 20pN, where minimal hairpin formation is expected. **(D)** Representative kymograph illustrating DNA replication under a tension of 20pN. The colocalization of fluorescent SSB with DNA polymerase is visible, suggesting that SSB facilitates the recruitment of DNA polymerase to bind on ssDNA.

**Figure S2.**
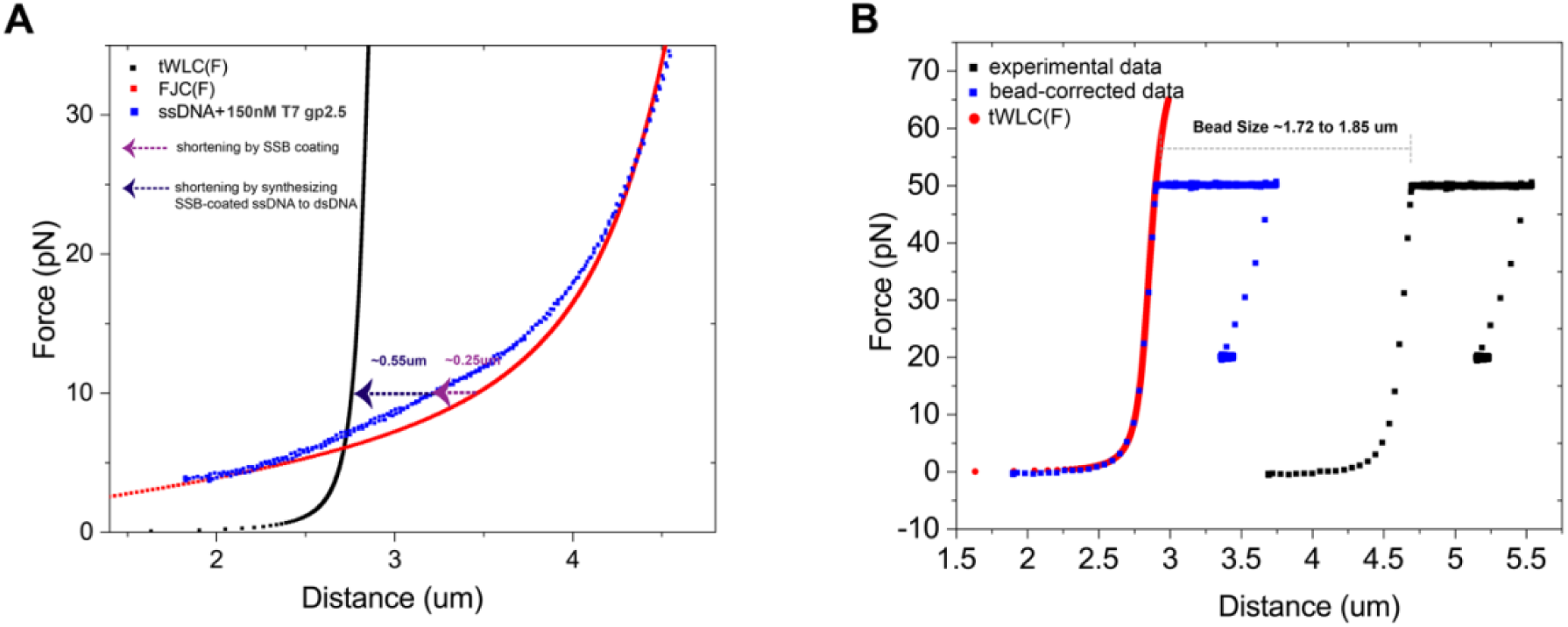
Estimation of the SSB binding calibration factor and bead size calibration. **(A)** Shows the correction factor, which accounts for changes in ssDNA length when it is fully coated by SSB proteins. The correction factor is estimated to be 0.246 µm under 10pN and 0.177 µm under 20pN with fully saturated gp2.5 protein. ‘FJC’ and ‘WLC’ refer to the Freely Jointed Chain and Worm-Like Chain models to describe ssDNA and dsDNA, respectively. **(B)** Bead size calibration for individual DNA molecules to account for variations in bead sizes. The experimental data curve (black line) is horizontally shifted to the left (blue line) to align with the theoretical WLC model. The difference in the positions of the black and blue lines represents the bead size, which in our experiments varies between 1.72 and 1.85 µm.

**Figure S3.**
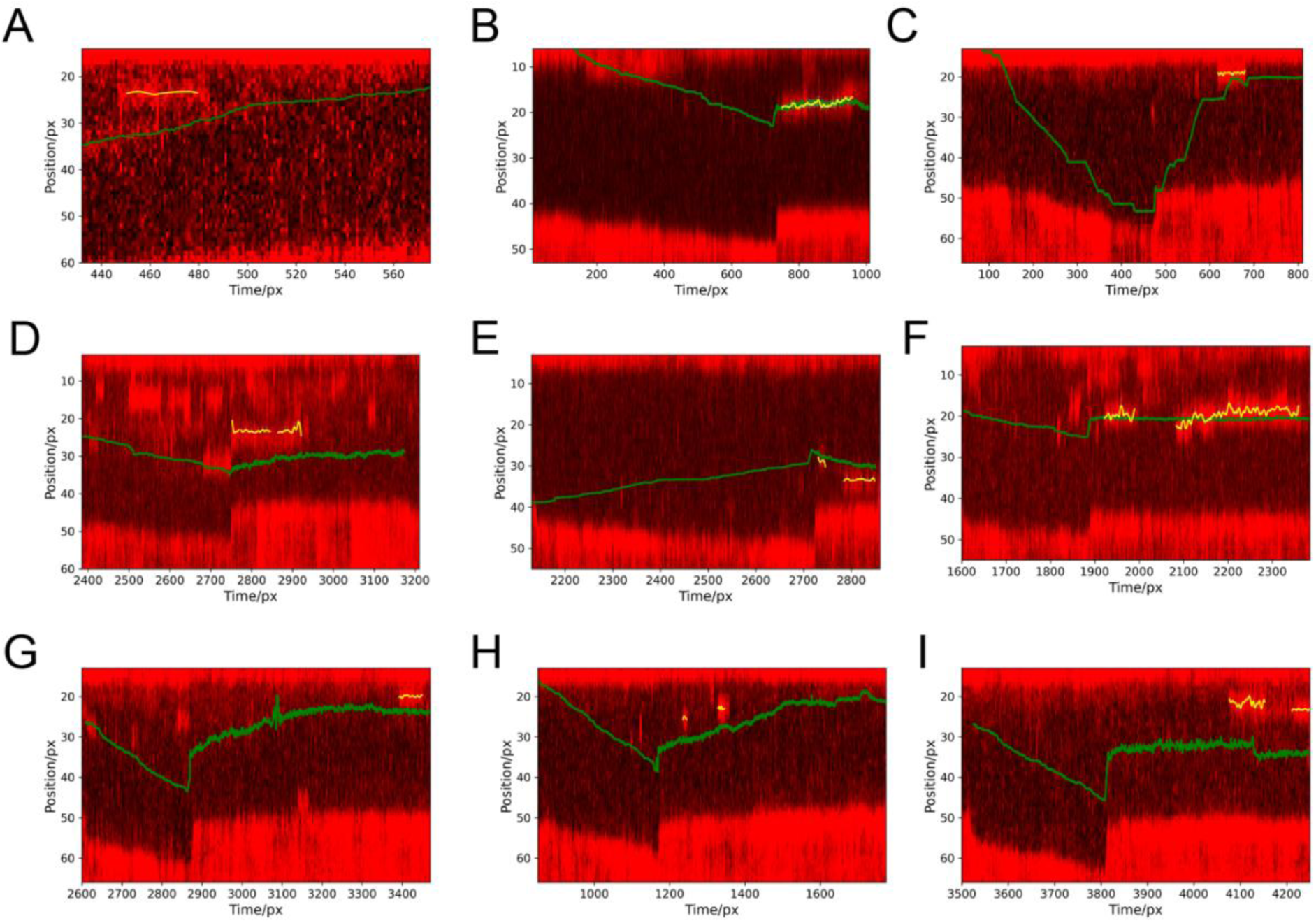
(A-I). A gallery of nine distinct DNA molecules showing the displacement of SSB by DNA polymerase. Each panel (A to I) displays a graphical representation of the position of the ssDNA/dsDNA junction as determined through independent optical tweezer measurements (represented by a green solid line), alongside the binding of fluorescently labeled SSB protein to ssDNA observed using fluorescence microscopy. The SSB binding data is extracted and represented by a yellow solid line, quantified via a trajectory analysis method. Only SSB binding events that last longer than approximately 2 seconds (3 pixels) are included in the analysis. The horizontal axis represents time, with an approximate scale of 0.7 seconds per pixel, while the vertical axis represents position, scaled at 75 nm per pixel. For details on the experimental conditions, see **Methods** section, and for an example of data analysis, see Figure 2.

**Figure S4.**
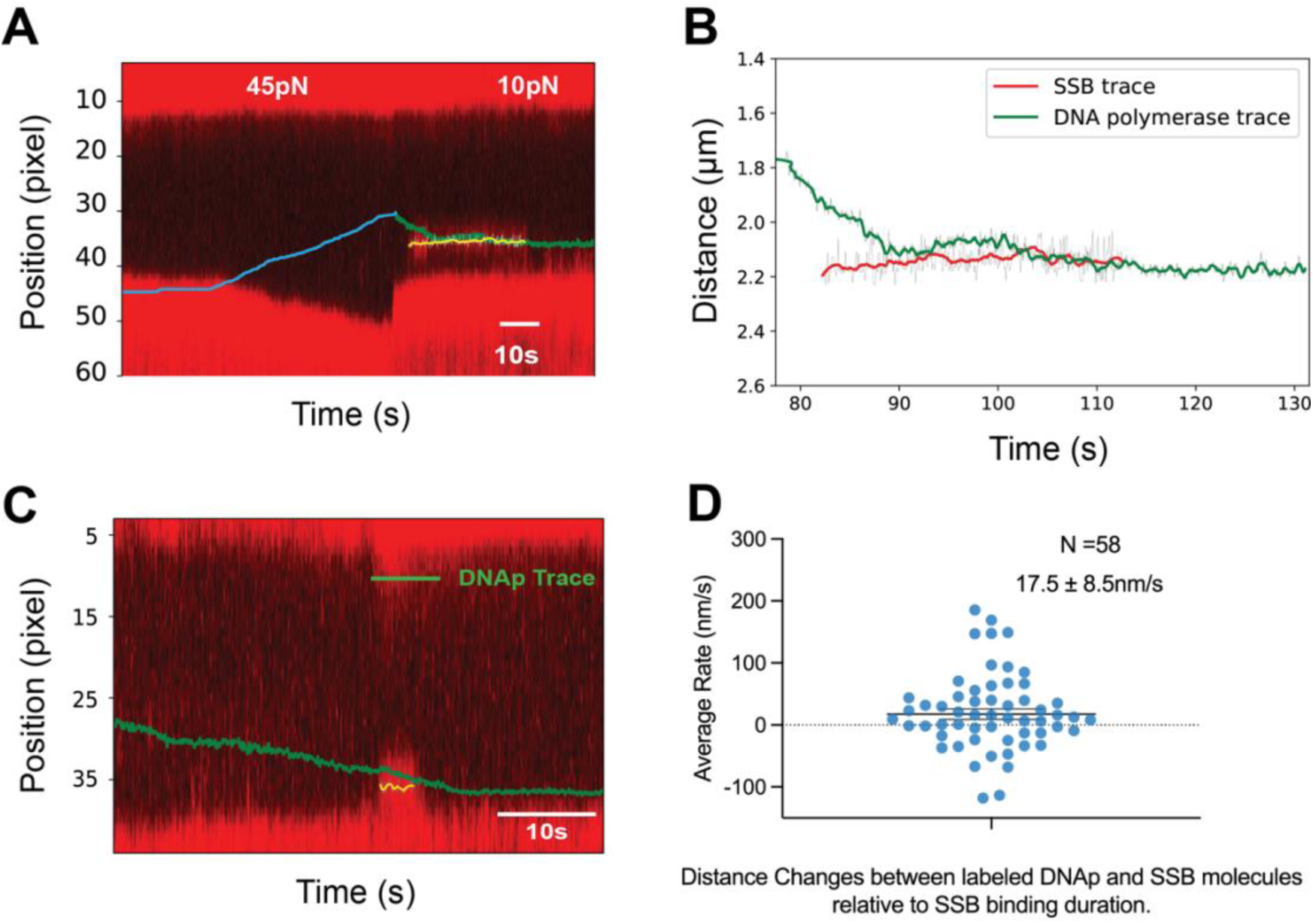
Tracking the real-time position of DNAp and SSB. **(A)** Fluorescence microscopy was used to directly observe interactions between DNA polymerase and SSB. The DNAp position (green) was tracked in real-time using optical tweezer measurements that exploit the elasticity differences between ssDNA and dsDNA (**Methods**). Fluorescently labeled SSB binding to ssDNA was visualized, and SSB trajectories (yellow) were extracted for further analysis using a trajectory analysis method (**Methods**). The data is from the same molecule as Figure 2D but includes both exonuclease and polymerization activities. **(B)** Fluorescent trajectories from (A) are synchronized with DNAp trajectories computed from ssDNA/dsDNA junction positions. The green and red traces represent the positions of fluorescently labeled DNAp and SSB over time, respectively. A Savitzky-Golay filter is applied to denoise the raw traces (shown in grey) for clarity. Only the polymerization process is displayed for clarity. **(C)** Representative traces showing the displacement of fluorescently labeled SSB by DNAp under 10 pN force (N= 2 out of 25 DNA molecules). Time scale: 10 s; position scale: 75 nm per pixel. **(D)** Net replication rate of DNA polymerase toward SSB during 58 individual binding events, calculated from the overall distance change between DNAp and SSB during the event’s binding lifetime. The average rate of DNAp movement toward SSB is 17.5 ± 8.5 nm/s (mean ± SEM), corresponding to approximately 200 bp/s, similar to the net DNA replication rate in **Figure S1B**. The broad distribution may result from intermittent pausing and proofreading during individual events.

**Figure S5.**
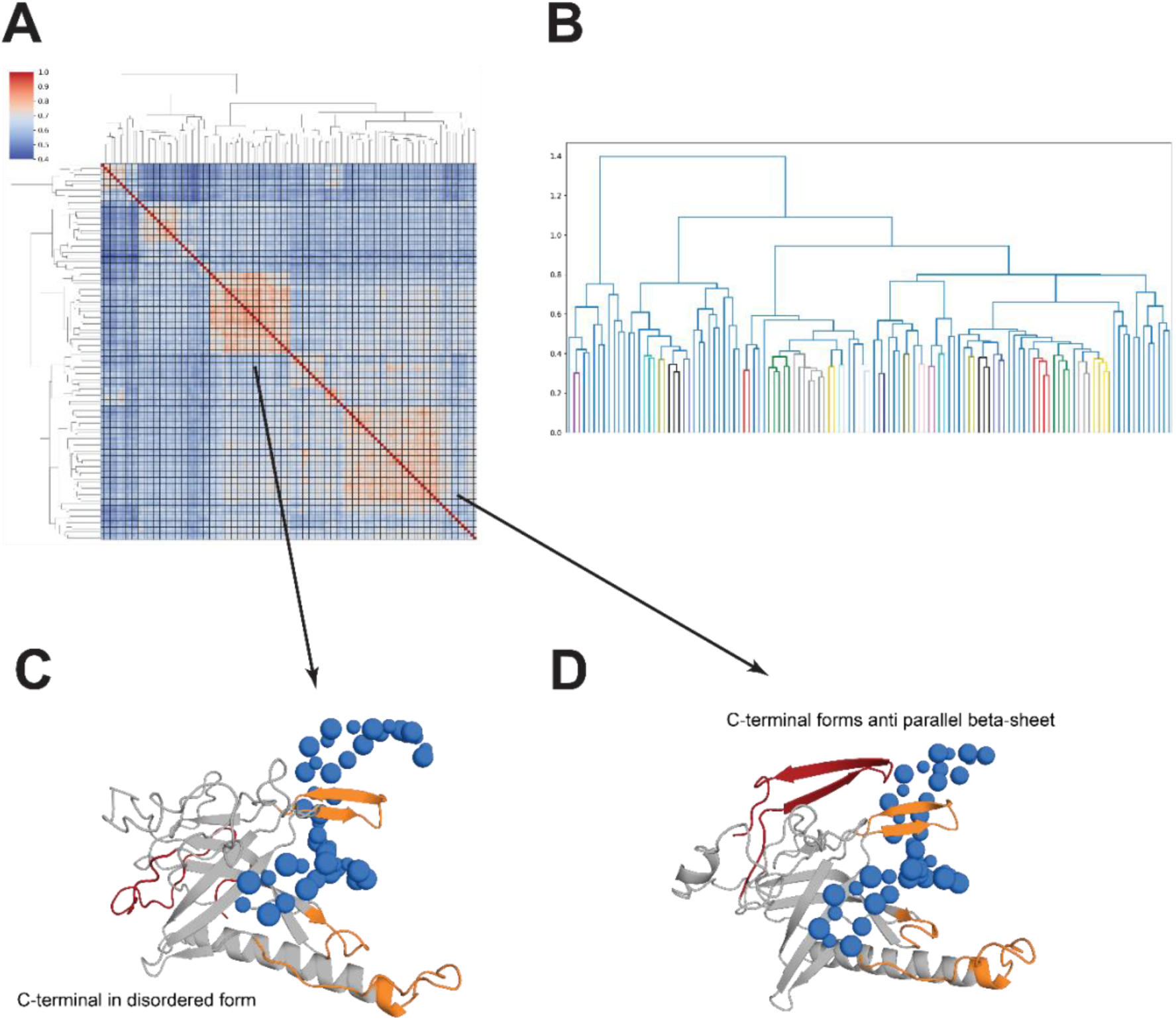
(A-D). The clustering analysis of the bound state of SSB-ssDNA complex. **(A)** The clustermap was computed by the mutual Qw value of each pair of protein that reside in the bound state of Figure 3D. The color indicates the range of Qw value, 1 (red) means the identical structure while 0 (blue) means totally dissimilar. **(B)** The dendrogram shows the cluster of these extracted structures. The center of each small cluster is shown in different colors while the remaining branches are color in blue. **(C)** The representative structure of one of the clusters in (A). This structure has a disordered C-terminal of SSB. **(D)** The representative structure of one of the clusters in (A). This structure has a C-terminal with anti-parallel beta-sheet secondary structure.

**Figure S6.**
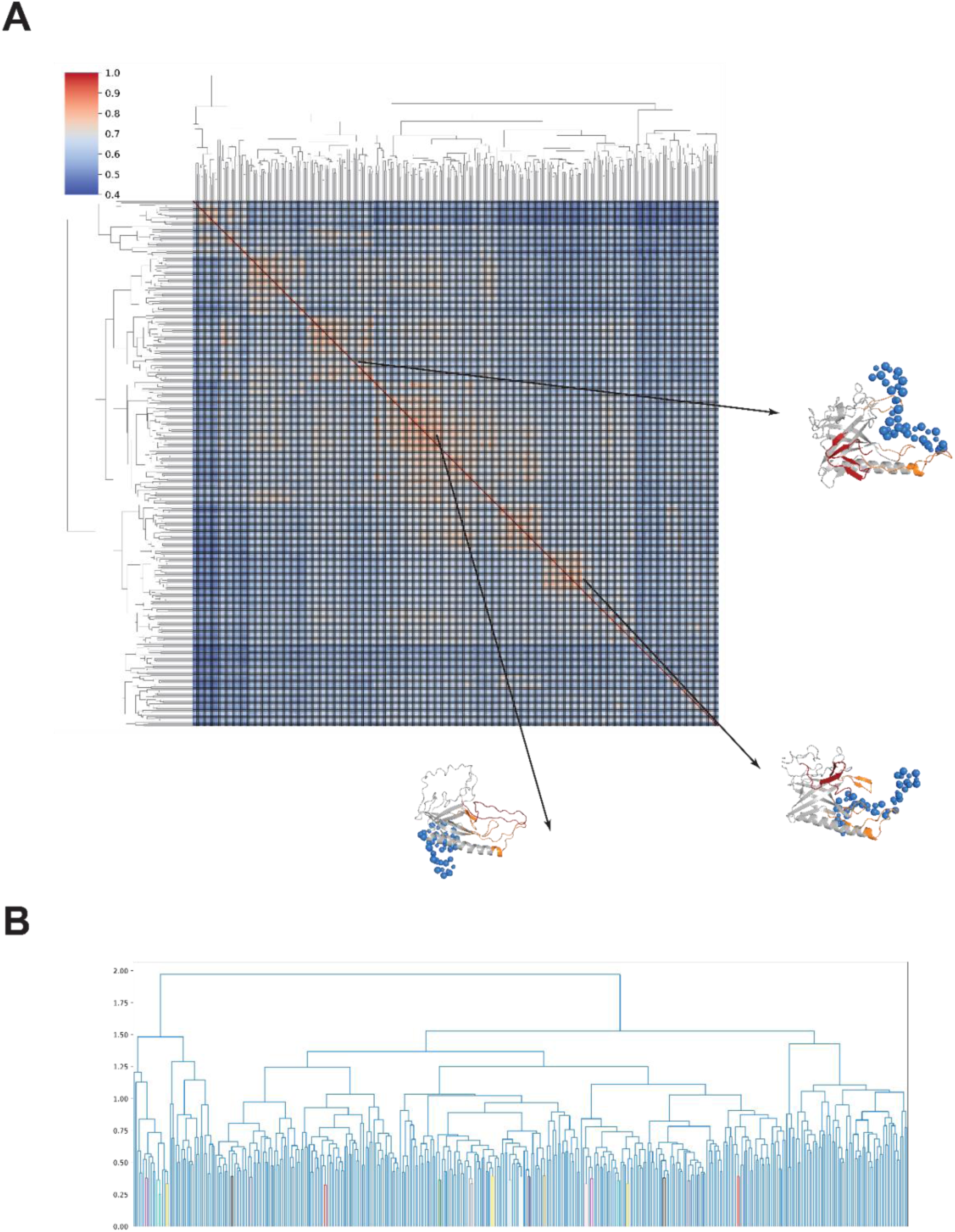
The clustering analysis of the intermediate state of SSB-ssDNA complex. **(A)** The clustermap was computed by the mutual Qw value of each pair of protein that reside in the intermediate state of Figure 3D. The color indicates the range of Qw value, 1 (red) means the identical structure while 0 (blue) means totally dissimilar. The representative structure was picked from the center of each cluster. **(B)** The dendrogram shows the cluster of these extracted structures. The center of each small cluster is shown in different colors while the remaining branches are color in blue.

**Figure S7.**
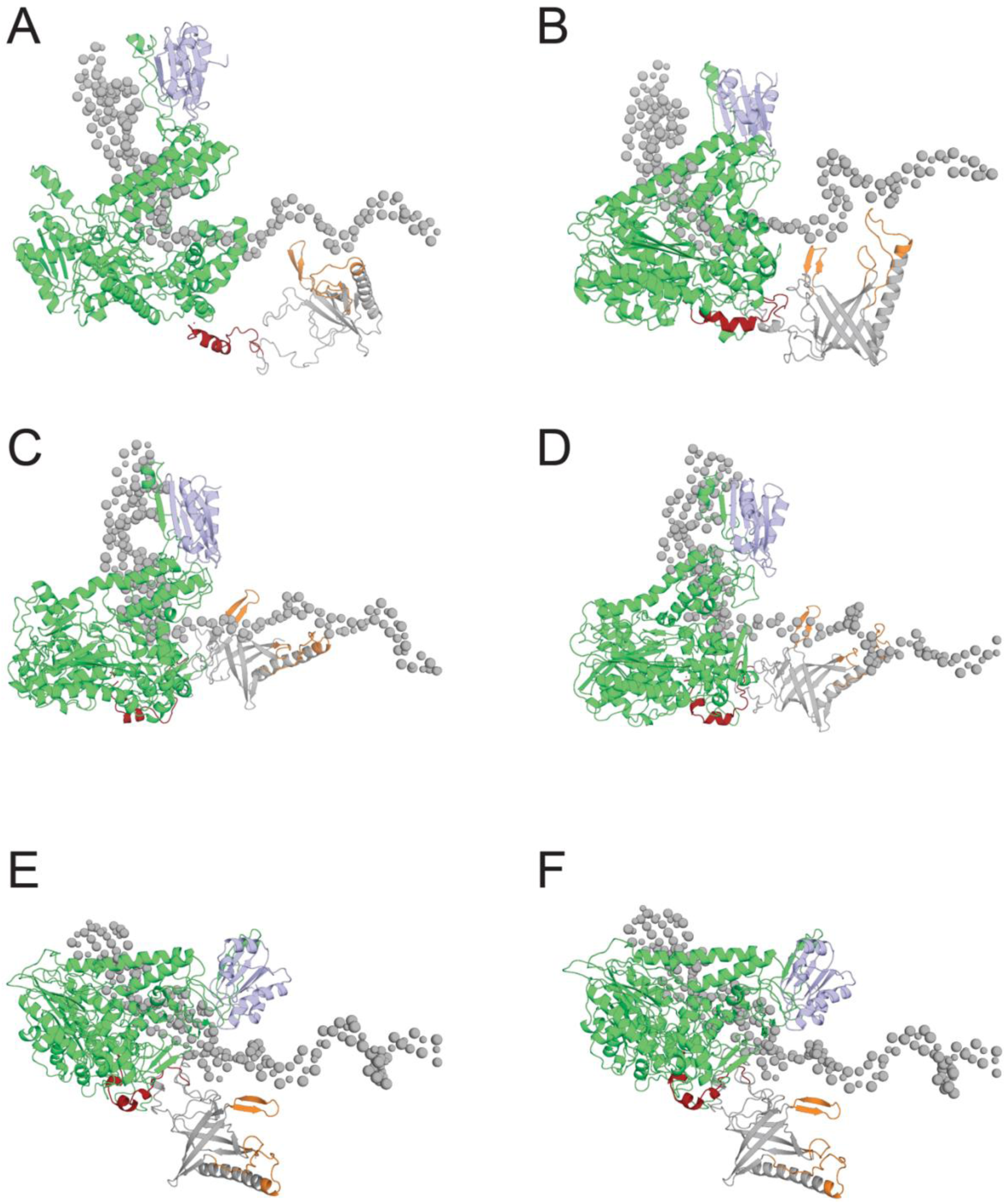
Representative snapshots of T7 SSB-DNAp complex trajectory. **(A)** The snapshots of initial state in the trajectory. At the beginning, the SSB sits in the side of the ssDNA. **(B)** The snapshots of bound state in the trajectory. The orange colored positive charged groove of SSB can find the correct binding position when interact with ssDNA. **(C)** The snapshots of the trajectory shows the interaction between SSB and ssDNA while maintain the contact with DNAp in one direction. **(D)** The snapshots of the trajectory shows the interaction between SSB and ssDNA while maintain the contact with DNAp in another direction. **(E)** The snapshots of the dissociated state in the trajectory shows the SSB has started to pull out from ssDNA. **(F)** The snapshots of the dissociated state in the trajectory shows SSB has completely removed from ssDNA.

**Figure S8.**
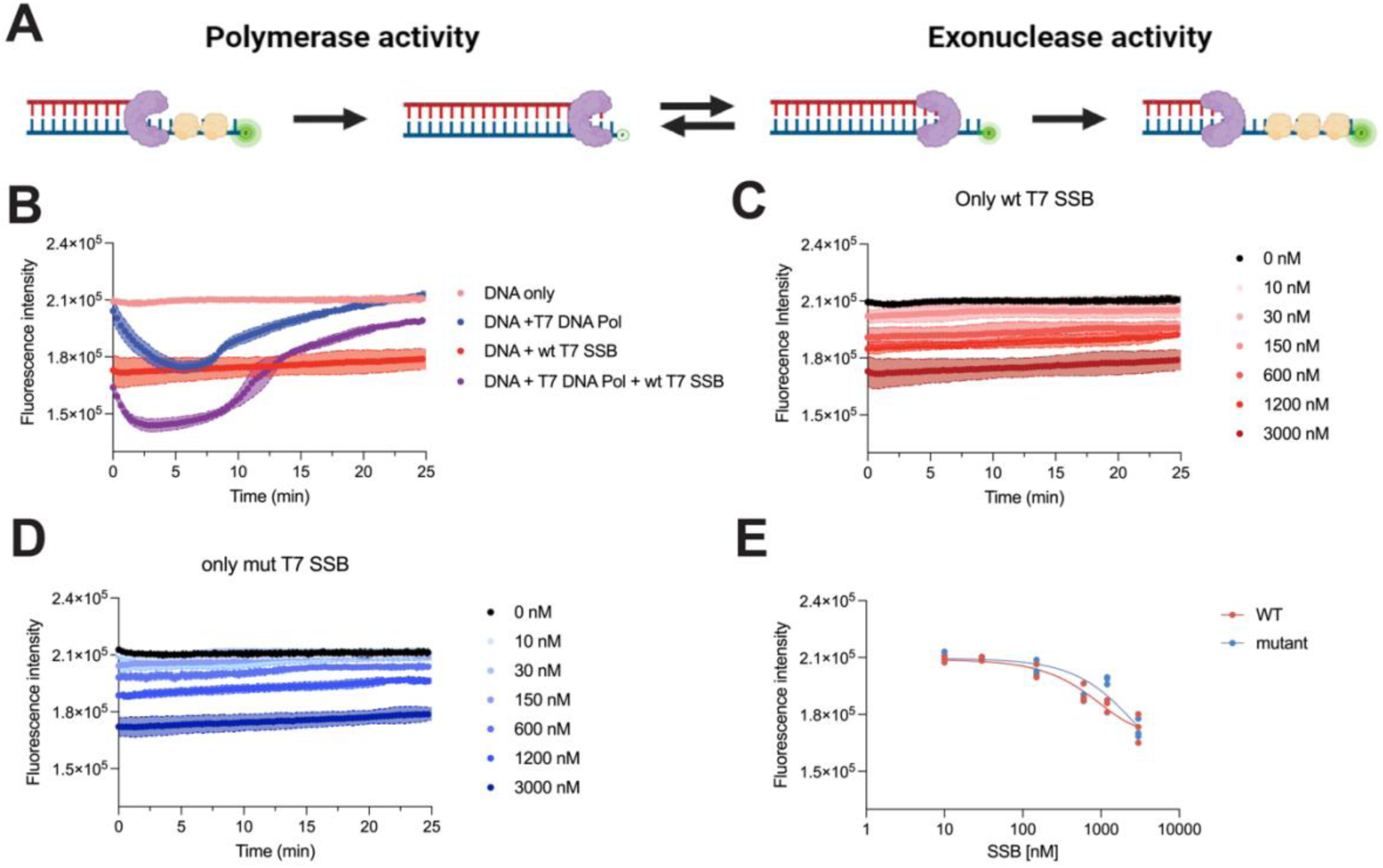
Real-time DNA primer extension assay investigating the interaction between DNA polymerase and SSB. **(A)** Schematic representation of the real-time assay. A 5′-end fluorophore-labeled template is quenched upon primer extension, allowing monitoring of DNA synthesis kinetics. Fluorescence signal decreases during polymerase activity and increases during exonuclease activity. Purple: DNA polymerase; Yellow: SSB. **(B)** Representative real-time polymerase assay traces showing average fluorescence signals for 70 nM DNA alone (light red), DNA with 30 nM DNA polymerase (blue), DNA with 3000 nM wild-type T7 SSB (red), and DNA with both 30 nM DNA polymerase and 3000 nM wt T7 SSB (purple). Initial fluorescence drop indicates primer extension. SSB presence accelerates polymerase activity. As the primer reaches full extension, polymerase and exonuclease activities reach equilibrium. Upon nucleotide depletion, exonuclease activity dominates, increasing the signal. **(C)** and **(D)** Control experiments showing real-time fluorescence signals with increasing concentrations (0, 10, 30, 150, 600, 1200, and 3000 nM) of wild-type SSB **(C)** or mutant SSB **(D)** with 70 nM DNA, without polymerase. Higher SSB concentrations correlate with decreased overall fluorescence. Notably, the signal remains stable over the experimental timeframe (∼30 min), enabling quantification of SSB effects on DNA polymerase. **(D)** Initial fluorescence signal plotted against increasing concentrations of wild-type (red) or mutant (blue) SSB. Both SSB variants show similar fluorescence reduction patterns, with effects noticeable from approximately 150 nM SSB concentration.

**Table S1.** Experimentally derived binding kinetics of SSB to ssDNA under varying template tension. Comparison of binding parameters between wild-type gp2.5 and gp2.5-ΔC21. The average binding free energy, on-rate (*kon*) per binding site, and off-rate (*koff*) are presented. *koff* values were obtained by fitting a triple exponential function, with k1 considered as *koff* due to the order of magnitude difference between k1, k2, and k3. On-rates and off-rates are sourced from **Supplementary Table 1** of our recent publication(***20***).

| Template Tension | SSB Type | $k_{on} (M^{-1} s^{-1} \text{ Site}^{-1})$ | $k_{off} (s^{-1})$ | $K_d = K_{off}/K_{on} (M \cdot \text{Site})$ | $\Delta G = RT \ln(K_d) (kJ/mol)$ | $\Delta G = RT \ln(K_d) (kcal/mol)$ |
| --- | --- | --- | --- | --- | --- | --- |
| 3pN | wt gp2.5 | $1.38E4 \pm 2.94E3$ | $4.98E0 \pm 6.01E-1$ | $3.61E-04$ | -1,98E+01 | -4,73E+00 |
| | gp2.5-ΔC26 | $5.11E3 \pm 7.28E1$ | $4.23 E0 \pm 6.57E-1$ | $8,28E-04$ | -1,77E+01 | -4,23E+00 |
| 6pN | wt gp2.5 | $1.30E4 \pm 5.63E2$ | $5.14E0 \pm 5.60E-2$ | $3,95E-04$ | -1,95E+01 | -4,67E+00 |
| | gp2.5-ΔC26 | $5.88E3 \pm 8.20E2$ | $4.33E0 \pm 6.81E-1$ | $7,36E-04$ | -1,80E+01 | -4,30E+00 |
| 12pN | wt gp2.5 | $1.07E4 \pm 2.49E3$ | $2.42E0 \pm 3.38E-2$ | $2,26E-04$ | -2,09E+01 | -5,00E+00 |
| | gp2.5-ΔC26 | $1.57E4 \pm 6.78E2$ | $4.81E0 \pm 1.11E-1$ | $3,06E-04$ | -2,02E+01 | -4,82E+00 |
| 18pN | wt gp2.5 | $5.59E3 \pm 4.35E0$ | $3.23E0 \pm 3.76E-3$ | $5,78E-04$ | -1,86E+01 | -4,44E+00 |
| | gp2.5-ΔC26 | $9.13E3 \pm 8.26E2$ | $5.02E0 \pm 1.62E-1$ | $5,50E-04$ | -1,87E+01 | -4,47E+00 |

**Movie S1. Morphing movie of the structures in the intermediate state.**

[view with this link]

A morph movie illustrates the potential moving direction from the representative structure picked from **Figure S5**. It’s worth to note the detailed path is unrealistic as the movie was generated using the simple morph function by Pymol. However, one can still observe the process that C-terminal attack the positive binding groove and repel the ssDNA from its original position.

**Movie S2. The unbias simulation of DNAp-SSB.**

[view with this link]

**Movie S3. The unbias simulation trajectory of DNAp-SSBΔ21C.**

[view with this link]

